# Lung tissue organization shapes influenza spread and immune control

**DOI:** 10.64898/2026.11.19.767682

**Authors:** Rodolfo Blanco-Rodriguez, Benjamin Whipple, Eugenia Yeboah, Reagan R. Johnson, Firas Maayah, Qasem Alharbi, Alec Smith, Yumna Kurdi, Tanya A. Miura, Esteban A. Hernandez-Vargas

## Abstract

Type I interferon (IFN) is a central defense against influenza A virus, yet its effectiveness depends on lung tissue organization, often simplified in mechanistic models. We investigated four IFN mechanisms—blocking infection, inducing a refractory state, suppressing virion production, and amplifying IFN production—using a homogeneous reaction–diffusion model and a spatial agent-based model reconstructed from murine lung histology. The latter represented approximately 233,000 detected cells in a single section or 4.1 million across a reconstructed volume. Models were calibrated against PR8 viral titers and IFN-*β* measurements, with spatial models additionally constrained by day-2 lesion extent. In the homogeneous model, bulk kinetics could not distinguish IFN mechanisms, and tissue sparing arose from widespread refractory conversion rather than spatial containment. In histology-derived networks, tissue organization slowed viral spread relative to a matched regular lattice, reducing cumulative infection from 98% to 55% under identical parameters. Under spatial constraints, the tested innate mechanisms could not reproduce both lesion development and viral clearance without CTL-mediated killing. Refractory conversion exhibited a sharp threshold: strong protection prevented infection establishment, whereas weaker protection reduced severity without reproducing clearance. Infection outcomes also depended on CTL expansion, inoculum size, and local tissue density. Despite differences in connectivity and fitted parameters, independently calibrated two- and three-dimensional representations supported the same mechanistic conclusions. Together, these findings show that tissue network organization shapes immune control and that spatial observations distinguish mechanisms appearing equivalent from bulk kinetics alone.

**SIGNIFICANCE STATEMENT:** Influenza spreads through the lung in scattered infection foci, yet many models represent tissue as a uniform environment. We reconstructed a cellular network from histological sections of an infected mouse lung, using the positions of approximately four million detected cells to simulate the movement of virus and interferon. We found that tissue organization slows viral spread, providing more time for killer T cells to control infection. Although strong interferon protection can prevent infection from becoming established, it cannot reproduce the observed infection dynamics without T-cell-mediated clearance. Infection severity also depends on where infection begins, how strongly cells respond to interferon, and how quickly adaptive immunity develops. These findings show why realistic tissue structure matters for understanding influenza infection and immune control.

---

**I**NFLUENZA A virus (IAV) does not fill the mouse lung; it colonizes it. Infection appears as multiple expanding foci separated by parenchyma that remains untouched, and large regions are still uninfected when viral load peaks (1–4). Whether such an infection remains focal or becomes extensive damage is decided in the first days, by the interplay between local cell-to-cell transmission, the geometry of the alveolar walls, and the reach of immune effectors (5, 6). Type I interferons (IFN-*α/β*) act first in this competition. Secreted by infected cells, IFN induces an antiviral state in neighboring cells, can suppress virion output from cells already infected, and amplifies its own production through IRF7-dependent feedback (7–13). In cell culture the paracrine antiviral state is established within hours of virion production and can arrest the spread of infection altogether (14). Which of these actions restricts spread through tissue, and whether any of them suffices before adaptive immunity arrives, remain open.

Mathematical models have carried most of the weight in addressing these questions, and they have done so at two separate spatial scales. Target-cell-limited ordinary differential equation models treat the lung as one well-mixed container and have been remarkably successful at describing viral kinetics (12, 15, 16), at the price of discarding space entirely. Explicitly spatial models—partial differential equations (PDE), agent-based models (ABM), cellular automata, and hybrids (Table S1)—recover local spread but are almost always solved on idealized domains (17–33). Recent work has linked local focus expansion to long-range seeding, identifying seeding as the main driver of early exponential growth, with limited IFN restriction for well-adapted viruses (34). However, this framework does not account for the spatial organization of the tissue. Here, we investigate how realistic lung tissue architecture influences IFN-mediated viral control.

Two features of lung tissue make its spatial organization particularly relevant. First, the alveolar architecture is highly heterogeneous, with epithelial cells lining thin walls separated by airspaces, interrupted by airways and vessels, and exhibiting substantial variations in local cell density and susceptibility to infection (35). Such heterogeneity may constrain viral spread through narrow, tortuous paths while also limiting the reach of IFN-mediated protection. Second, IAV models are typically calibrated against aggregate measurements that capture infection dynamics but provide little information about spatial spread, making it difficult to distinguish mechanisms with similar temporal predictions but different spatial effects (5).

We compared four IFN mechanisms—blocking of infection (BI), refractory conversion (RF), blocking of virion production (BP), and positive feedback on IFN production (PF)— across four spatial representations (Fig. 1): a homogeneous reaction–diffusion model and a common agent-based model (ABM) implemented on a regular lattice, a histological section of an infected mouse lung, and a three-dimensional reconstruction from twenty serial sections (4.1 million cells). All models were calibrated to per-animal PR8 viral titers and IFN-*β* fold changes (36), with ABMs additionally constrained by immunohistochemical measurements of infection extent at day 2.

**Fig. 1.**
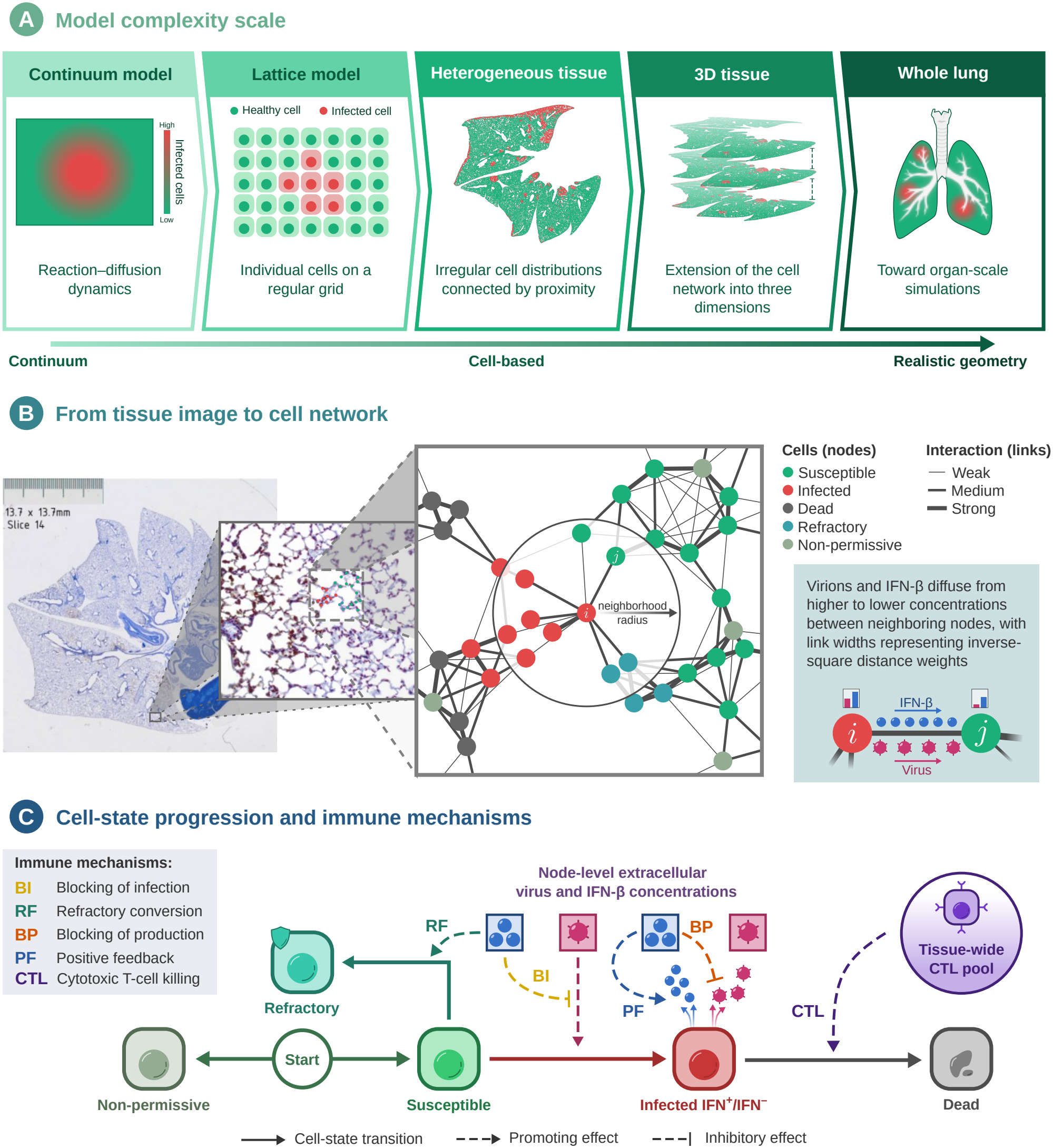
Tissue representations, cell network, and model mechanisms. (*A*) Spatial representations ranging from a continuum and regular lattice to a histological section, a reconstructed three-dimensional volume, and the whole organ. (*B*) Cell-to-network conversion: cells identified in histological images become nodes connected to nearby neighbors, with distance-dependent weights governing virion and IFN-*β* diffusion. The circle highlights the neighborhood of node *i*, showing only its connections for clarity; connections between neighbors are omitted within the circle. (*C*) Cell states and immune mechanisms: blocking of infection (BI) and refractory conversion (RF) act on susceptible cells, while blocking of production (BP) and positive feedback (PF) act on infected cells. An IFN-driven, tissue-wide cytotoxic T-lymphocyte (CTL) response eliminates infected cells throughout the tissue.

We find that bulk kinetics cannot distinguish IFN mechanisms in homogeneous tissue, where tissue sparing produces uniform dilution rather than spatial barriers. In reconstructed tissue, innate mechanisms alone fail to reproduce the observed dynamics, requiring an IFN-recruited systemic cytotoxic T-lymphocyte (CTL) response for infection clearance. Tissue network topology slows viral spread, providing time for this response, while IFN mechanisms primarily modulate tissue damage rather than contain infection. Finally, two- and three-dimensional tissue representations yield distinct parameter estimates that do not transfer between them, highlighting the importance of representation-specific spatial calibration.

## Results

### Bulk kinetics identify no IFN mechanism on homogeneous tissue

We first evaluated whether bulk infection kinetics could distinguish IFN mechanisms in a homogeneous reaction–diffusion model. Fifteen combinations of the four IFN mechanisms were independently calibrated to experimental viral titers and IFN-*β* fold changes (36). The positive feedback (PF)-only combination was excluded because it has no effect on infection dynamics in the absence of other IFN mechanisms. All models reproduced the observed kinetics with nearly identical likelihoods, leaving the model without IFN mechanisms as the most parsimonious (Tables S2 and S3; Fig. S1). However, the fits did not constrain the spatial extent of infection: most predicted infection of nearly the entire domain, with viral decline driven primarily by target-cell depletion.

Because influenza infection in the lung is focal and much of the tissue remains spared (1), we repeated the calibration requiring at least 80% of cells to remain uninfected by day 10. This constraint favored refractory conversion but did not improve the ability of bulk kinetics to distinguish mechanisms. Instead, IFN-mediated protection was distributed throughout the domain, while infection reached every location by day 3 (Fig. S1*E,F*). Thus, the homogeneous model achieved tissue sparing through widespread refractory conversion rather than spatial containment. These results motivate examining how realistic tissue architecture alters viral spread and IFN-mediated protection using the agent-based model. Full model specifications, calibration procedures, and additional results are provided in *Supplementary Material*, section 1.

### Real tissue network topology slows viral spread

To move beyond the homogeneous continuum model toward a more realistic representation of tissue structure, we developed TissueFlu, a GPU-based agent-based model (ABM) that represents individual cells at arbitrary spatial coordinates, without requiring a regular lattice. Through image processing of twenty serial histological sections of a PR8-infected mouse lung, we identified the positions of approximately 4.1 million cells, with infected cells detected by hemagglutinin immunostaining. The first section (denoted z0; 233,479 identified cells) and the complete stack provided two- and three-dimensional tissue representations, respectively. These cells were represented as nodes in a distance-weighted network (Fig. 1*B*). Each cell is initially assigned a susceptible or non-permissive state. Local extracellular virus promotes infection of susceptible cells, while infected cells produce virions, may produce IFN, and eventually die. Local IFN mediates four mechanisms: blocking of infection (BI), refractory conversion (RF), blocking of virion production (BP), and positive feedback on IFN production (PF). A systemic cytotoxic T-lymphocyte (CTL) response, modeled as being recruited by tissue-wide IFN, eliminates infected cells (Fig. 1*C*; Materials and Methods).

To isolate the effect of tissue network topology, we compared z0 with a regular hexagonal lattice matched in cell number, mean connectivity, and mean coupling strength. Despite comparable average connectivity, the real tissue network exhibited greater heterogeneity, lower reachability, and reduced effective dimensionality (1.71 versus 1.90; Fig. 2*A–C*; Table S6). These structural differences substantially altered infection dynamics under identical model parameters.On the regular lattice, infection spread as circular foci that rapidly coalesced, whereas on the histology-derived tissue network, spread remained irregular and patchy (Fig. 2*D*; Movies S1 and S2). By day 2, the fraction of infected or dead cells reached 33% on the lattice, compared with 8% on real tissue and 13 ± 5% experimentally. Viral load peaked earlier on the lattice (day 2.8 versus 4.3), with greater tissue damage (98% versus 55% of permissive cells; Fig. 2*E,F*).

**Fig. 2.**
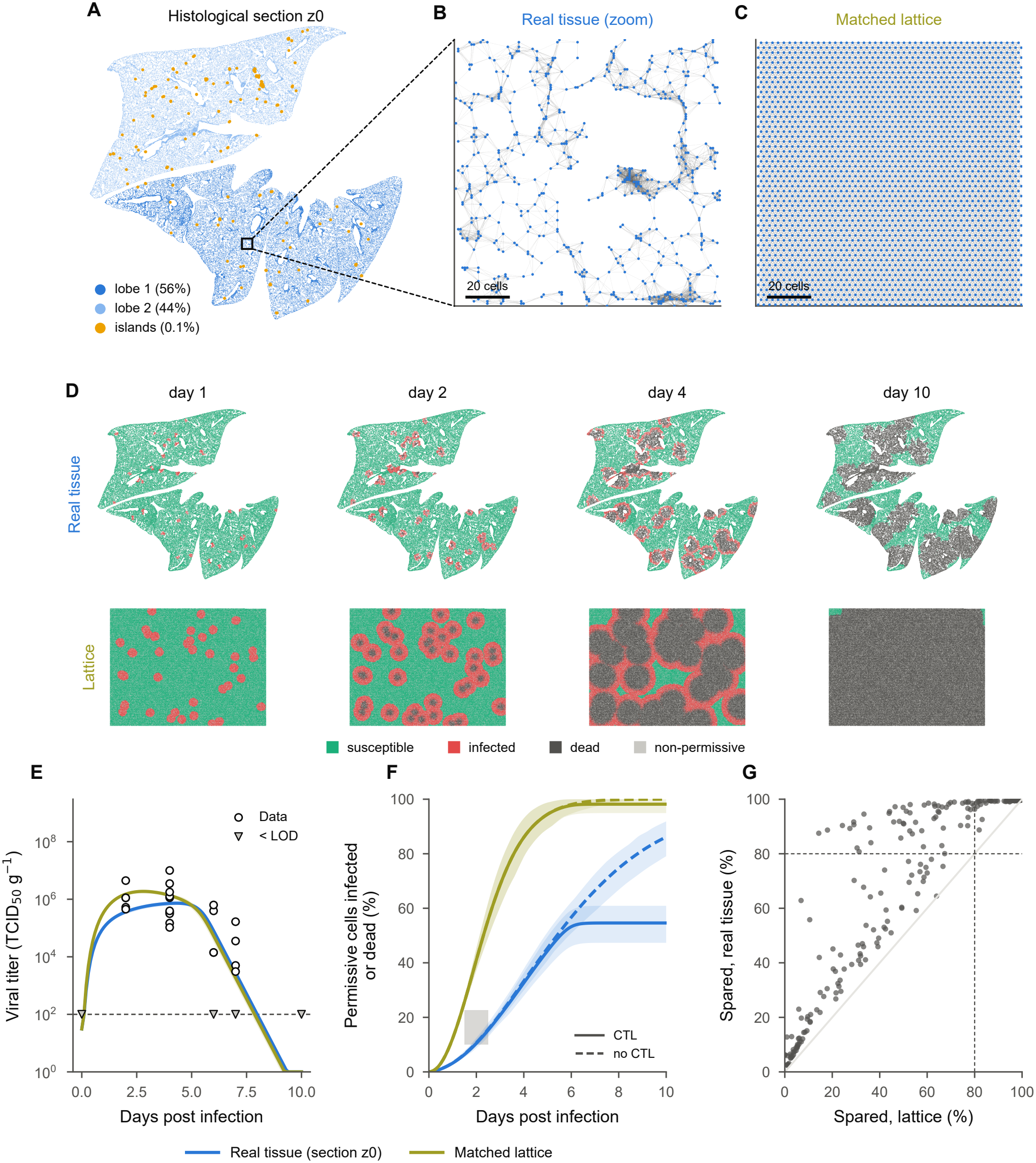
Real tissue network topology slows viral spread relative to a homogeneous lattice with matched mean coupling. (*A*) Section z0 colored by connected components of the cell network: two major lobes (~56% and ~44% of cells) and small islands (less than 0.1%; enlarged for visibility). The square indicates the region enlarged in *B*. (*B,C*) Cell positions (dots) and network connections (lines) in a 120 *×* 120 region of the real tissue (*B*) and a hexagonal lattice matched in cell number, mean connectivity, and coupling strength (*C*). (*D*) Cell states at days 1, 2, 4, and 10 under identical parameters (best z0 model: CTL without IFN mechanisms; one replicate). (*E–F*) Blue, real tissue; olive, matched lattice. (*E*) Viral titers for the same model on both networks (50 replicates; offsets from the z0 calibration). (*F*) Fraction of permissive cells infected or dead with (solid) and without CTLs (dashed); shaded bands indicate the 95% replicate range. Gray region indicates the measured day-2 lesion extent. (*G*) Fraction of permissive cells spared at day 10 across 400 combinations of IFN mechanisms and transport parameters without CTLs (two replicates each); dashed lines indicate 80% sparing.

Without CTLs, infection on the lattice rapidly spread throughout the domain, leading to viral clearance through depletion of permissive cells rather than spatial containment. Real tissue followed the same trend but progressed more slowly, with approximately 90% of permissive cells infected or dead by day 10 and virus still detectable (Fig. 2*F*; Table S7). Thus, the rapid clearance observed on homogeneous tissue resulted from target-cell exhaustion, whereas realistic tissue network topology delayed this process.

To test whether stronger IFN responses could contain infection without CTLs, we sampled 400 combinations of IFN mechanism strengths and transport parameters on both networks. Real tissue spared more cells than the lattice in 395 cases, but containment occurred only when infection remained nearly abortive: none of the 299 contained simulations reproduced the observed lesion extent (Fig. 2*G*; Fig. S6). On the lattice, containment compatible with the observed lesion extent was possible only through widespread refractory conversion, as in the continuum model, and yielded poor fits. Together, these results show that realistic tissue network topology slows viral spread but does not enable IFN alone to contain infection at the observed scale.

### CTL-mediated clearance is required to reproduce infection dynamics

We next evaluated which immune mechanisms were required to reproduce infection dynamics on real tissue by calibrating all fifteen IFN-mechanism combinations with and without cytotoxic T lymphocytes (CTLs). In the two-dimensional section (z0), the best innate-only model required three mechanisms (BI, RF, BP) but failed to reproduce the observed viral kinetics, peaking too early and declining too slowly (Fig. 3*A*). In contrast, including CTLs reproduced the viral peak and subsequent decline below the detection limit by days 8–10, with no additional IFN mechanism favored by model selection (Table S4). Both models captured the IFN-*β* dynamics, but CTL-mediated clearance occurred without the extensive refractory conversion associated with innate-only control (Fig. 3*B,C*).

**Fig. 3.**
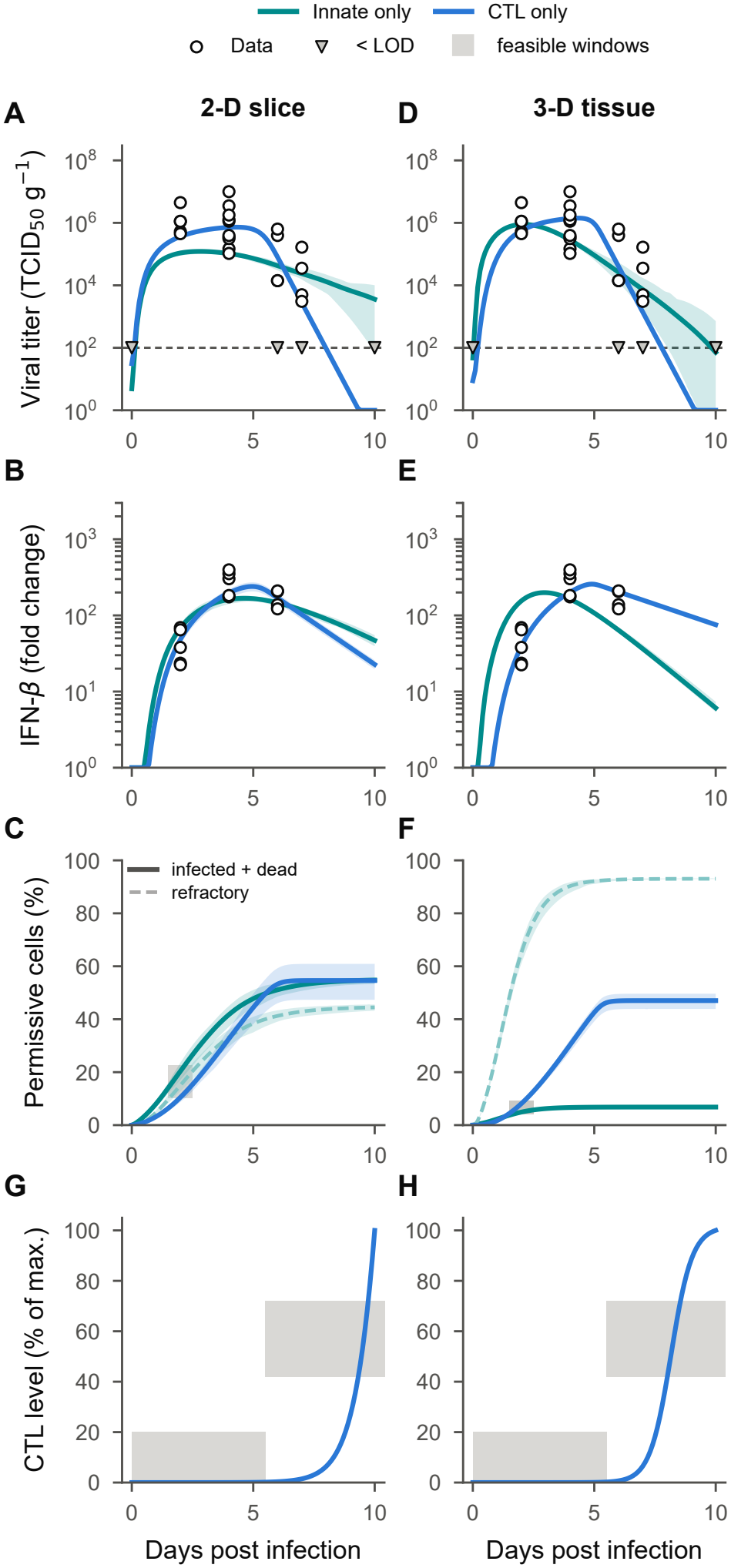
CTL-mediated clearance is required to reproduce infection dynamics on reconstructed lung tissue. (*A–C*) Two-dimensional section (z0): viral titer (*A*), IFN-*β* fold change (*B*), and fraction of permissive cells infected or dead (solid) or refractory (dashed) (*C*). Curves show the best models without CTLs (dark cyan) and with CTLs but no direct IFN antiviral mechanisms (blue). (*D–F*) Corresponding results for the three-dimensional reconstruction, comparing the best innate-only (dark cyan) and CTL-only (blue) models. (*G,H*) Systemic CTL population, expressed as a percentage of its maximum, for the calibrated CTL-only models in 2-D (*G*) and 3-D (*H*). Gray regions in *C* and *F* indicate feasible day-2 lesion extents derived from histological measurements with adopted uncertainty margins; those in *G* and *H* indicate prescribed feasible windows for early and late CTL population levels. Lines show median trajectories, and shaded bands indicate the 95% replicate range.

The three-dimensional reconstruction reinforced the importance of CTLs despite differences in network topology and spatial calibration. The independently calibrated CTL-only model reproduced viral and IFN-*β* kinetics, the measured day-2 lesion extent, and viral clearance by day 8 (Fig. 3*D– F*; Table S4). Without CTLs, no tested parameter set simultaneously satisfied these constraints. The best innate-only model fitted substantially worse (ΔAICc ≈ 102) and relied on widespread refractory conversion, reaching approximately 93% of permissive cells by day 10, while failing to reproduce the observed viral kinetics and clearance (Fig. 3*D–F*). The contrasting infection outcomes further illustrate the importance of CTL-mediated clearance: the innate-only model limited cumulative infection to approximately 7% of permissive cells through extensive refractory conversion, whereas the CTL-only model allowed approximately 47% to become infected or die before clearance.

The calibrated CTL populations exhibited delayed expansion in both tissue representations (Fig. 3*G,H*). CTL levels remained low during early infection before increasing rapidly, with the three-dimensional model showing an earlier response than the two-dimensional model. Both trajectories satisfied the prescribed feasible windows for early and late CTL population levels (37). The corresponding probability of CTL-mediated killing reached half-maximal values on days 7.4 and 6.2 in 2-D and 3-D, respectively, reflecting the nonlinear dependence of killing probability on CTL abundance rather than the time at which the CTL population reached half of its maximum. Together, these results indicate that the tested IFN-mediated mechanisms alone were insufficient to reproduce the observed infection dynamics, whereas CTL-mediated killing provided the clearance mechanism in both tissue representations.

Despite the stochastic nature of TissueFlu, replicate simulations showed limited variability in aggregate infection dynamics, as reflected by the narrow 95% replicate intervals around the median trajectories (Fig. 3*A–F*). This variability was particularly small once infection became established, when the large number of interacting cells reduced the relative influence of stochastic fluctuations. Thus, stochastic cell-level dynamics produced nearly deterministic population-level behavior under the conditions examined.

### Calibrated parameters across tissue representations

To assess how spatial representation influences parameter estimation, we compared the CTL-only models independently calibrated on the two-dimensional section and three-dimensional reconstruction (Table 1). Both models share the same biological rules, with infection kinetics parame-ters fixed to reduce the number of free parameters. Calibration against viral titers, IFN-*β* measurements, and day-2 lesion extent therefore allowed us to examine how differences in tissue connectivity and dimensionality affect the inferred immune-response parameters.

**Table 1.**
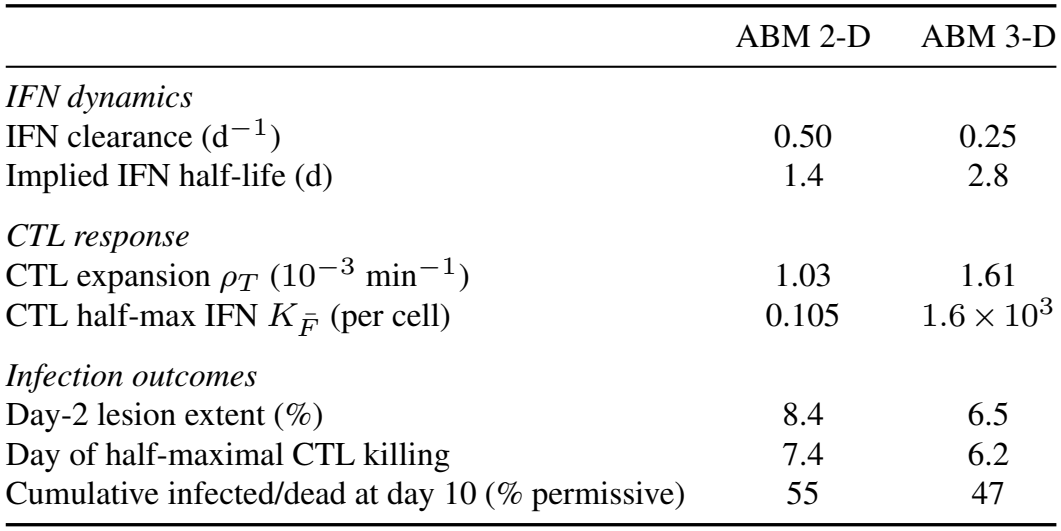
Comparison of calibrated immune-response parameters and predicted infection outcomes between the two-dimensional (2-D) and three-dimensional (3-D) agent-based models. Both models include CTL-mediated killing without direct IFN antiviral mechanisms. Full parameter lists, bounds, and units are provided in Table S5.

|  | ABM 2-D | ABM 3-D |
|---|---|---|
| *IFN dynamics* |  |  |
| IFN clearance (d<sup>−1</sup>) | 0.50 | 0.25 |
| Implied IFN half-life (d) | 1.4 | 2.8 |
| *CTL response* |  |  |
| CTL expansion *ρ<sub>T</sub>* (10<sup>−3</sup> min<sup>−1</sup>) | 1.03 | 1.61 |
| CTL half-max IFN $K_{\bar{F}}$ (per cell) | 0.105 | 1.6 × 10<sup>3</sup> |
| *Infection outcomes* |  |  |
| Day-2 lesion extent (%) | 8.4 | 6.5 |
| Day of half-maximal CTL killing | 7.4 | 6.2 |
| Cumulative infected/dead at day 10 (% permissive) | 55 | 47 |

The two representations yielded different estimates of IFN turnover and CTL dynamics. The fitted IFN clearance rate was 0.50 d^−1^ in 2-D and 0.25 d^−1^ in 3-D, corresponding to half-lives of 1.4 and 2.8 days, respectively. The CTL expansion rate increased from 1.03 × 10^−3^ to 1.61 × 10^−3^ min^−1^, while the half-maximal IFN threshold for CTL expansion differed substantially (0.105 versus 1.6 × 10^3^ IFN units per cell). Because this threshold depends on the IFN concentrations generated by each model, the difference should not be interpreted directly as a change in biological IFN sensitivity. Despite these parameter differences, both models reproduced the observed viral and IFN kinetics and predicted day-2 lesion extents of 8.4% in 2-D and 6.5% in 3-D. Half-maximal CTL killing occurred on days 7.4 and 6.2, respectively, and by day 10, the cumulative fraction of permissive cells infected or dead reached 55% and 47%. Thus, independently calibrated tissue representations supported the same requirement for CTL-mediated clearance and produced broadly similar infection outcomes.

### IFN sensitivity, CTL timing, and inoculum location shape infection severity

We next examined how IFN mechanisms modulate cumulative infection by independently varying their strengths in the calibrated z0 model (Fig. 4*A*). Blocking of infection (BI) and virion production (BP) modestly reduced cumulative infection from 55% to 38–39% of permissive cells at their strongest levels, while positive feedback (PF) alone had negligible effects because IFN-dependent CTL recruitment was already saturated. In contrast, refractory conversion (RF) showed a sharp threshold-like response: cumulative infection dropped from 44% to 15% over one strength increment and below 1% at the next. This transition corresponded to an increase in the local refractory conversion probability from 1.4% to 5.4% per minute, with nearly abortive infection at 20%. Although strong RF could contain infection without CTLs, the resulting lesions remained far below the experimentally observed extent (Fig. S4). Combining PF with intermediate RF further reduced cumulative infection from 44% to 21%, revealing a cooperative effect between these mechanisms (Fig. S5).

**Fig. 4.**
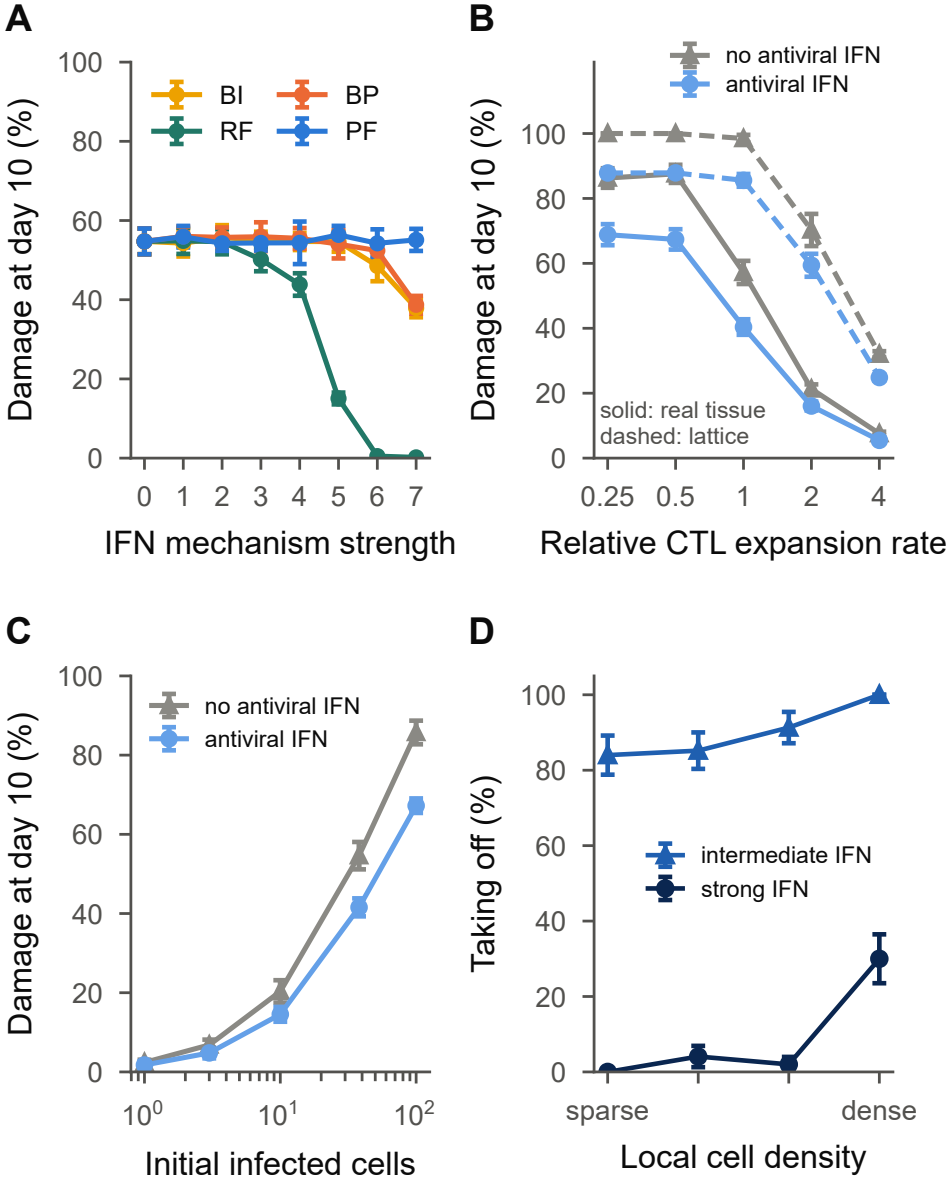
IFN sensitivity, CTL expansion, and inoculum conditions shape infection severity. (*A*) Cumulative infection (permissive cells infected or dead) at day 10 as each IFN mechanism is added independently to the calibrated z0 model (CTL with-out IFN mechanisms) at seven strength levels (0, inactive; 7, strongest). Mean *±* s.d. of 10 replicates. (*B*) Cumulative infection at day 10 across CTL expansion rates relative to the calibrated value (1), comparing real tissue and the matched lattice, with and without antiviral IFN protection. (*C*) Cumulative infection at day 10 as a function of the number of initially infected cells, without (gray) and with (blue) antiviral IFN protection; mean *±* s.d. of 20 replicates. (*D*) Fraction of single-cell inocula establishing infection across quartiles of local cell density, measured within 30 cell diameters of the inoculation site, under intermediate and strong antiviral IFN protection (200 inocula per IFN condition in total; error bars, binomial s.e.).

CTL expansion timing also strongly influenced infection severity (Fig. 4*B*). On real tissue, doubling the calibrated expansion rate reduced cumulative infection from 57% to 21%, whereas halving it increased infection to approximately 86%. On the matched lattice, faster viral spread left less time for CTL-mediated control. Thus, tissue network topology delayed infection and widened the window for an effective adaptive response. An intermediate antiviral IFN response further reduced cumulative infection across CTL expansion rates but did not substantially shift this timing dependence. Inoculum size and location introduced additional variability in infection outcomes. Increasing the number of initially infected cells from one to 38 and 100 raised cumulative infection from approximately 2% to 55% and 86%, respectively, consistent with the formation of additional infection foci (Fig. 4*C*). Local cell density also influenced infection establishment: under strong antiviral IFN protection, none of the inocula in the sparsest quarter of tissue locations established infection, compared with 30% in the densest quarter (Fig. 4*D*). Under intermediate protection, these proportions increased to 84% and 100%, respectively.

Together, these results show that infection severity depends on IFN sensitivity, CTL expansion timing, and the size and location of the initial inoculum. Tissue network topology shapes these effects by controlling the rate of viral spread and the time available for immune control.

### A single tissue section supports mechanistic inference when spatially calibrated

The neighborhood radius *r* determines which cells can exchange virus and IFN, with markedly different consequences across tissue representations (Fig. 5; Table S6). In individual sections, the largest connected component expands sharply between *r* = 6 and *r* = 8, but the network remains divided into two major components even at *r* = 20 (Fig. 5*A*). These sections also contain persistent spatial gaps: at *r* = 9–12, persistent homology identified 335–372 holes per 10^4^ cells, including large gaps that persist across a broad range of neighborhood radii (Fig. 5*D*). Across four sections sampled at different depths, connectivity, effective dimension, reachability, and hole-size distributions were broadly similar, supporting the use of a representative section.

**Fig. 5.**
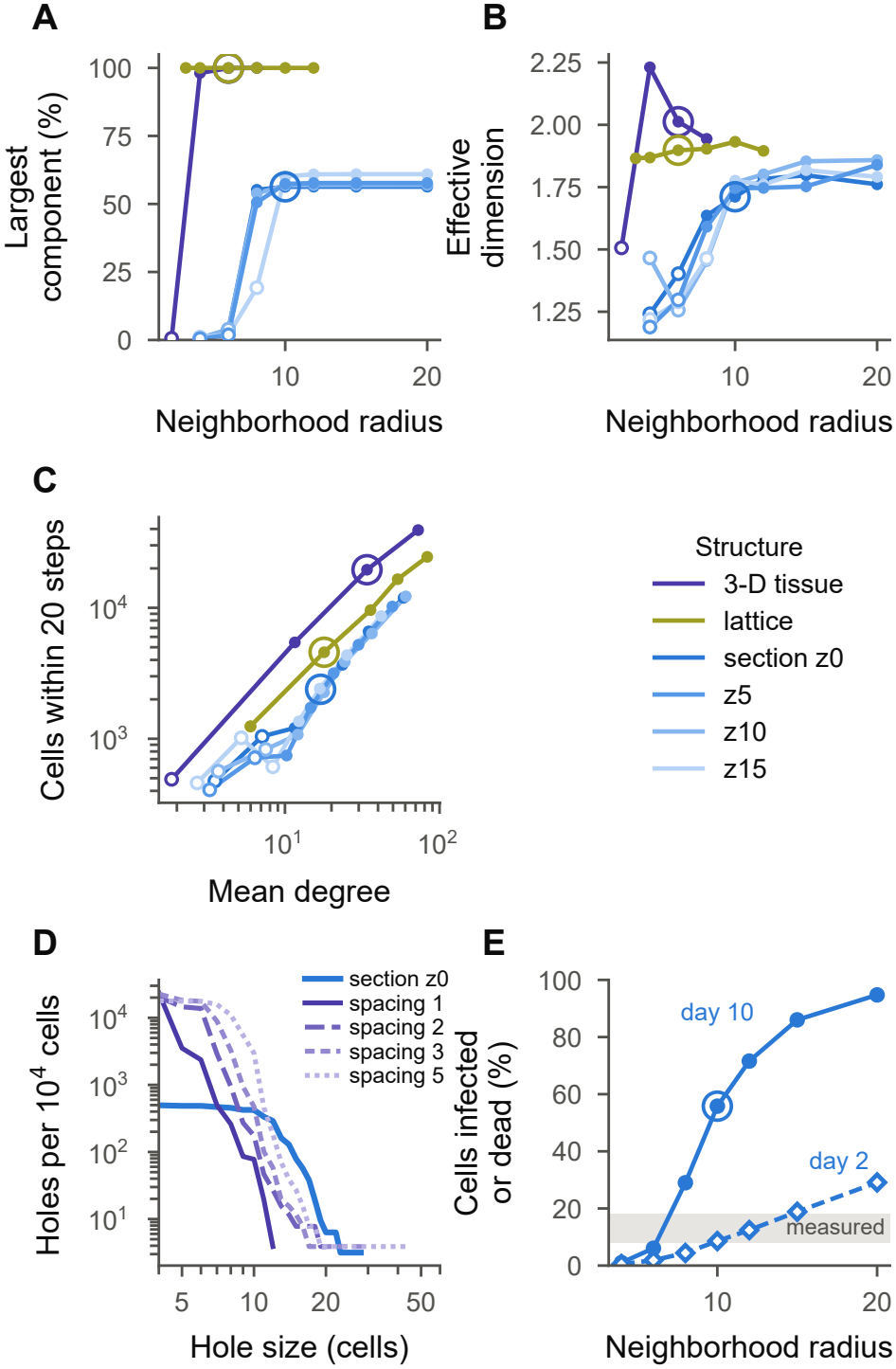
Single tissue sections show consistent network properties, whereas three-dimensional connectivity depends on section spacing. (*A–C*) Network properties of four tissue sections, the reconstructed three-dimensional volume, and the matched lattice: fraction of cells in the largest connected component (*A*), effective dimension as a function of neighborhood radius (*B*), and number of cells reachable within 20 network steps as a function of mean degree (*C*). Circled points indicate radii used in simulations; hollow markers indicate radii at which the largest connected component contains less than half of the cells. (*D*) Number of holes per 10^4^ cells requiring at least a given radius to be bridged, for a region of section z0 (blue) and the 20-section volume reconstructed with inter-section spacings of 1, 2, 3, or 5 units (purple). (*E*) Fraction of permissive cells infected or dead at days 10 (solid) and 2 (dashed) across neighborhood radii, using the calibrated z0 model with diffusivities rescaled to preserve mean coupling strength. Gray band indicates the experimentally measured day-2 lesion extent; circled point marks the calibrated radius.

In contrast, the stacked three-dimensional volume forms a single connected component at *r* = 4, has a higher effective dimension, and allows cells to reach substantially more neighbors within 20 network steps (Fig. 5*A–C*). At equal neighborhood radius, reachability was approximately eightfold higher than in a section and remained 3–4.5-fold higher when matched by mean degree. Nevertheless, the effective dimension of the volume remained close to 2, suggesting that infection spread is still largely constrained to a two-dimensional network at this spatial resolution. Thus, although stacking sections increases connectivity, it does not introduce a fully three-dimensional spreading structure, and individual sections may capture much of the spatial organization relevant to infection propagation. Accordingly, the neighborhood radius required to reproduce the observed day-2 lesion extent differed between representations: *r* = 10 for the section and *r* = 6 for the volume.

However, the connectivity of the reconstructed volume depends on the assumed spacing between histological sections. Increasing the spacing from one to five units enlarged network gaps and reduced the mean degree at the calibrated radius from 35 to 10 (Table S8). The number of holes spanning at least 12 units increased from 4 to 183 per 10^4^ cells, approaching the 291 observed in a single section (Fig. 5*D*). Because the physical separation between sampled sections is uncertain, the inferred three-dimensional connectivity is sensitive to a reconstruction assumption that cannot be independently constrained by these data.

Network connectivity also strongly influenced infection severity (Fig. 5*E*). In the calibrated z0 model, increasing the neighborhood radius from 4 to 20 raised cumulative infection at day 10 from 1% to 95% of permissive cells, closely following the growth of the largest connected component. This dependence persisted when virus and IFN diffusivities were rescaled to preserve mean coupling strength (Fig. S7). Thus, increasing the number of reachable cells, rather than simply increasing average coupling strength, was the dominant effect of expanding the interaction neighborhood in these simulations.

Finally, we tested whether calibrated parameters could be transferred across tissue representations. Applying the z0 parameters to three other sections produced broadly similar outcomes, although cumulative infection was lower in the least connected section (Fig. S8). In contrast, transferring the same parameters to the three-dimensional volume, after rescaling diffusivities, preserved a good fit to bulk kinetics but overestimated the day-2 lesion extent (15% versus 5.3% observed) and increased cumulative infection to 83% (Fig. S7*B,D*). Thus, agreement with bulk measurements alone did not guarantee accurate spatial predictions. When calibrated independently against their respective spatial observations, both representations supported the same mechanistic conclusions: CTL-mediated clearance was necessary to reproduce the observed infection dynamics, whereas sufficiently strong refractory conversion could prevent infection establishment but not reproduce the observed lesion extent. These findings support the use of a single tissue section for the mechanistic questions examined here, while highlighting that calibrated parameters cannot be transferred directly between two- and three-dimensional representations.

## Discussion

Mechanistic models of influenza infection are often used to infer immune mechanisms from viral titers. Our results show that such inference is strongly influenced by how tissue structure is represented. In a homogeneous continuum, fifteen combinations of four IFN mechanisms reproduced the bulk kinetics with comparable accuracy, yet implied markedly different levels of tissue involvement. Even imposing substantial tissue sparing did not produce spatial containment: infection reached the entire domain, while IFN-mediated refractory conversion left susceptible cells distributed throughout it. Thus, sparing cells was not equivalent to restricting the spatial spread of infection. This distinction extends the limitations of target-cell-limited explanations of viral clearance (5): bulk kinetics alone cannot establish whether declining viral loads reflect immune-mediated clearance, target-cell depletion, or spatial containment.

Previous modeling work has demonstrated that spatially localized autocrine and paracrine IFN signaling can arrest viral spread by inducing antiviral states in neighboring cells (12). Using ODE, PDE, and cellular automaton models, that study showed how spatial structure can enhance IFN-mediated protection. Our work extends this comparison to histology-derived lung networks, with models additionally constrained by measured viral and IFN kinetics and early lesion extent. Although sufficiently strong IFN responses could prevent infection establishment, the tested innate mechanisms could not reproduce both lesion development and viral clearance without CTL-mediated killing. Moreover, replacing a regular lattice with measured tissue connectivity substantially altered infection outcomes. These findings do not contradict the protective potential of paracrine IFN signaling but demonstrate that its inferred sufficiency depends on tissue organization and experimental constraints.

Beyond its influence on viral spread, spatial modeling can help distinguish the contributions of different immune mechanisms. For example, a model of herpes simplex virus-2 infection incorporating experimentally measured tissue-resident T-cell densities identified cytokine-mediated antiviral effects as a dominant control mechanism (33). In our study, the measured day-2 lesion extent ruled out scenarios that reproduced viral and IFN kinetics but predicted inconsistent spatial infection patterns. Under these constraints, target-cell depletion and the tested innate mechanisms could not jointly reproduce lesion development and subsequent viral clearance, whereas models including CTL-mediated killing could. This is consistent with the established contribution of CD8^+^ T cells to influenza control (38, 39), although our model does not exclude other clearance mechanisms. Tissue organization also altered the competition between infection and adaptive immunity. Compared with a matched regular lattice, the heterogeneous tissue network reduced reachability and delayed viral spread (Fig. 2), leaving more cells uninfected as CTL activity increased. This delay widened the opportunity for CTL-mediated control, demonstrating that tissue connectivity influences not only infection progression but also the effectiveness of immune control.

IFN-mediated protection played distinct roles (Fig. 4*A*). Blocking infection and virion production reduced cumulative infection, whereas refractory conversion exhibited a sharp threshold: strong protection prevented infection establishment but also suppressed lesion development below the observed extent. Positive feedback enhanced protection mainly when combined with intermediate refractory conversion. Thus, IFN sensitivity influenced infection establishment and severity without necessarily determining viral clearance. In our model, tissue-wide IFN also stimulated CTL expansion, but this response was already saturated early under calibrated conditions, limiting the effect of further IFN amplification on CTL timing (Fig. 4*B*). This reflects the simplified coupling between innate and adaptive immunity rather than the full biological role of IFN in regulating adaptive responses.

Our results complement analytical work suggesting that IFN-mediated restriction may have limited effects on the early expansion of well-adapted respiratory viruses (34). By representing local infection on a measured cell network, we additionally identify how spatial organization changes the consequences of antiviral protection. In particular, the sharp response to refractory conversion suggests that modest differences in antiviral sensitivity could produce substantially different outcomes near the protection threshold. Infection establishment also depended on inoculum size and local cell density, with sparsely connected regions less likely to sustain infection under strong antiviral protection (Fig. 4*D*). These predictions are consistent with the broader importance of type I IFN in limiting influenza replication and disease, although its effects depend on viral strain and host background (40–42). Together, they suggest that local tissue organization and host antiviral sensitivity may jointly contribute to heterogeneity in infection outcomes.

The comparison between two- and three-dimensional representations highlights the importance of spatial calibration. Although the reconstructed volume increased connectivity and reachability, its effective dimension remained near 2 at the scales examined, and its connectivity depended strongly on the assumed inter-section spacing (Fig. 5). Transferring parameters from the section to the volume overestimated lesion extent despite reproducing bulk kinetics. However, when calibrated independently, both representations reproduced similar infection dynamics and supported the same requirement for CTL-mediated clearance (Fig. 3). Thus, a single histological section may provide sufficient spatial information for the mechanistic questions examined here, while increasing dimensionality does not necessarily improve inference without independently constrained tissue geometry.

Several limitations define the scope of these conclusions. The reconstructed tissue comes from one lung at a single time point, and uncertainty in section registration and spacing affects the inferred three-dimensional network (Table S8). Cells remain stationary, and the model does not explicitly represent airway transport, mucociliary clearance, immune-cell migration, or the diversity of pulmonary cell types. CTL activity is represented by a well-mixed compartment rather than explicit trafficking and antigen-specific interactions. Infection also spreads only through local network connections; long-range seeding between distant regions, which can generate spatially separated infection foci, is not represented (2, 3, 34). Combining local spread on measured tissue networks with long-range seeding would therefore provide a natural extension.

The IFN representation introduces further limitations. Whole-lung IFN-*β* transcript measurements constrain a single extracellular IFN field, although transcription, secretion, transport, and clearance occur on different timescales. The relatively slow IFN clearance inferred by the models may therefore reflect processes not explicitly represented, rather than extracellular protein stability alone. IFN production is heterogeneous in our model, with each infected cell assigned a fixed probability of producing IFN, consistent with single-cell studies showing that only a fraction of infected cells activate IFN production (43, 44). However, this probability was not independently calibrated, and additional cell-to-cell variability in IFN production was not represented. Similarly, the nonpermissive cell fraction was fixed, motivated by experimental evidence of heterogeneous cellular susceptibility to influenza infection (35). Virion release kinetics and infected-cell lifetime were also determined from reported literature rather than independently identified (45). Future measurements of lesion expansion at multiple time points, spatial IFN-stimulated gene expression, and infection dynamics under targeted immune perturbations could help distinguish the contributions of individual IFN mechanisms and better constrain their parameters.

Overall, our findings show that lung tissue organization shapes the competition between viral spread and immune control. By slowing local infection, heterogeneous connectivity changes the time available for adaptive immunity and the outcomes produced by IFN-mediated protection. Spatial measurements therefore provide constraints that bulk kinetics cannot, allowing mechanistic explanations of influenza infection to be evaluated against both temporal dynamics and tissue-level spread.

## Materials and Methods

### Experimental data

All models were calibrated against in vivo measurements of pulmonary PR8 viral load and IFN-*β* expression (36). Female BALB/c mice were inoculated intranasally with ~ 100 TCID_50_ of influenza A/Puerto Rico/8/1934 (H1N1); only the single-infection arm was used. Infectious viral load was quantified by TCID_50_ assay of whole-lung homogenate at days 2, 4, 6, 7, and 10 (five animals per day), with a detection limit *L* = 100 TCID_50_ g^−1^; titers at or below it were treated as left-censored, as were five day-0 observations representing the inoculum. IFN-*β* transcript was measured by RT-qPCR of whole-lung RNA, normalized to GAPDH and expressed as fold change relative to mock at days 2, 4, and 6. In total *n* = 50 observations enter the likelihood. The timing of the CTL response was constrained by two windows of the normalized T-cell level (37): 0–20% of maximum before day 5.5 and 42–72% be-tween days 5.5 and 13.5.

### Immunohistochemistry and reconstruction

Formalin-fixed lungs embedded in paraffin were cut into 5 µm sections and stained as described (36); IAV hemagglutinin was labeled with a polyclonal goat antibody (NR-3148, BEI Resources) and an HRP-conjugated anti-goat polymer (Vector Laboratories), visualized with DAB and counterstained with hematoxylin. Cell coordinates were extracted from whole-section images by combining high-resolution camera imaging with microscope-based calibration; binary masks of cell locations and infection status were generated with a custom image-processing workflow. Twenty serial sections were aligned by registration, giving 4,137,043 cells of which 217,123 (5.25%) were IAV-positive. Coordinates were scaled so that the minimum distance between cells is one unit, and consecutive sections were taken one unit apart.

### Reaction–diffusion model

We modeled infection on a homogeneous two-dimensional domain using a reaction– diffusion system tracking susceptible cells (*S*), infected cells (*I*), extracellular virus (*V*), and IFN (*F*). Four IFN mechanisms were considered: blocking infection (BI), refractory conversion of susceptible cells (RF), blocking virion production (BP), and positive feedback on IFN production (PF). Virus and IFN followed porous-medium diffusion, allowing finite-speed propagation, with IFN diffusivity set to 100 times that of virus. Simulations used a unit square with zero-flux boundaries. The governing equations, mechanism combinations, and constrained calibration procedure are detailed in *Supplementary Material*, section 1.

### Agent-based model

We developed TissueFlu, a stochastic agent-based model (ABM) that represents individual cells at arbitrary spatial coordinates. Each cell *i* is connected to neighboring cells within a radius *r*, with distance-dependent weights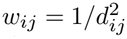. Extracellular virions and IFN are exchanged between connected cells through weighted diffusion, with local production and clearance. The model advances in discrete time steps of Δ*t* = 1 min, and cells are initially assigned as susceptible or nonpermissive with probability *p*_np_.

Cell-state transitions are stochastic and governed by Hill functions of local viral and IFN concentrations. Susceptible cells may become infected in response to extracellular virus or enter a refractory state through IFN-mediated protection (RF). IFN can also reduce the probability of infection (BI) and suppress virion production by infected cells (BP). Infected cells release virions according to single-cell production kinetics fitted to experimental data (45) and die after a stochastic lifetime with mean *τ*_*I*_. Upon infection, cells are assigned IFN-producing competence with probability *p*_+_; competent cells produce IFN through an intracellular dsRNA-dependent process that can be amplified by positive feedback (PF).

Adaptive immunity is represented by a single systemic cytotoxic T-lymphocyte (CTL) population, whose expansion follows a discrete-time logistic equation stimulated by the mean extracellular IFN concentration across the tissue. The probability of killing an infected cell is determined separately by a saturating Hill function of CTL abundance. This phenomenological formulation captures delayed systemic immune control without explicitly modeling CTL trafficking or spatially resolved interactions with infected cells.

Infection is initiated by depositing 100 virions at each of *N*_0_ selected cells. We used *N*_0_ = 38 for the two-dimensional section, corresponding to the number of IAV-positive clusters identified by HDBSCAN, and *N*_0_ = 100 for the reconstructed volume (Fig. S2). TissueFlu was implemented in CUDA C/C++, with one GPU thread per cell. On a single NVIDIA RTX A2000, a 10-day simulation requires approximately 3 s for a section and 90 s for the volume. Complete equations, transition probabilities, and parameter values are provided in *Supplementary Material*, section 2 and Table S5.

### Calibration and model selection

Models were calibrated against viral titers and IFN-*β* measurements using a censored-Gaussian negative log-likelihood with profiled log-scale offsets and pooled standard deviations (*σ*_*V*_ = 0.62, *σ*_*F*_ = 0.16). For the ABM, the objective was evaluated using the mean trajectories of replicates in which infection became established, with additional penalties for day-2 lesion extent and T-cell response timing (37). Model selection used the corrected Akaike information criterion, AICc = 2*J* +2*k* +2*k*(*k* +1)*/*(*n* −*k*− 1), where *J* is the negative log-likelihood, *k* the number of fitted parameters, and *n* = 50 the number of observations. Calibration combined Latin-hypercube sampling, trust-region refinement, and validation with 30–50 stochastic replicates (Fig. S3). The neighborhood radius and initial inoculum size (*N*_0_) were selected separately using a preliminary screen of day-2 lesion extent. Further details are provided in *Supplementary Material*, section 3.

### Network characterization and controls

For each tissue representation and neighborhood radius, we constructed the corresponding distance-weighted network and quantified its degree, coupling strength, connected components, clustering, reachability, and effective dimension. The latter was estimated from the scaling of the number of reachable cells with network distance over 3–20 steps. Spatial gaps were characterized using persistent homology of the distance-to-cell field, implemented with GUDHI (46). As a homogeneous spatial control, we constructed a regular hexagonal lattice of 233,479 cells, matched to the reference tissue section in mean degree and mean coupling strength. Network definitions, persistent-homology filtering, and lattice construction are detailed in *Supplementary Material*, section 3.

## DATA AVAILABILITY

Viral titer and IFN-*β* data are from ref. (36). The TissueFlu simulator is available at https://github.com/systemsmedicine/TissueFlu.git.

AI use disclosure: OpenAI’s ChatGPT (GPT-6) was used to assist with English language editing and improving manuscript clarity. Anthropic’s Claude (Opus 5.5) was used to assist with optimization of the custom agent-based model code and preparation of figures. All AI-assisted outputs were reviewed and verified by the authors, who take full responsibility for the scientific content, computational implementation, and results presented in this manuscript.

## AUTHOR CONTRIBUTIONS

R.B.-R., B.W., and E.H.-V. designed research; R.B.-R. and B.W. developed the methodology and software; E.Y. and T.A.M. prepared histological sections; F.M., Q.A., A.S., and Y.K. processed histological images; R.B.-R. and E.H.-V. wrote the original draft; R.B.-R., R.J., and E.H.-V. reviewed and edited the manuscript; and E.H.-V. supervised the research, administered the project, and acquired funding.

## ACKNOWLEDGEMENTS

Research reported in this publication was supported by the National Institute of General Medical Sciences of the National Institutes of Health under Award Numbers P30GM159567 and R01GM152736 (to E.H.-V.). The content is solely the responsibility of the authors and does not necessarily represent the official views of the National Institutes of Health. Simulations were run on the Lemhi high-performance computing cluster at the University of Idaho (47).

