## Supplementary Material for "Lung tissue organization shapes influenza spread and immune control"

Table S1 compares the two models used in this study with previously published spatial models of acute viral infection in epithelial tissues, focusing on their spatial geometry and calibration data. Sections 1–3 describe the models and tissue networks, Section 4 presents additional results supporting the main text, and Sections 5–7 contain the supplementary tables, figures, and movies.

#### 1. The reaction–diffusion model

**Governing equations.** The continuum model represents the epithelium as a homogeneous two-dimensional domain without an adaptive immune compartment. Susceptible and infected cell densities are denoted by $S(\mathbf{x},t)$ and $I(\mathbf{x},t)$, respectively, and extracellular virion and IFN-$\beta$ concentrations by $V(\mathbf{x},t)$ and $F(\mathbf{x},t)$. All variables are defined on the unit square $\Omega = [0,1]^2$. With all four IFN mechanisms active, the governing equations are

$$\partial_t S = -\beta \frac{k_{\mathrm{IF}}}{k_{\mathrm{IF}}+F} VS - k_R \frac{F}{K_R+F} S, \quad \textbf{(S1)}$$

$$\partial_t I = \beta \frac{k_{\mathrm{IF}}}{k_{\mathrm{IF}}+F} VS - \delta_I I, \quad \textbf{(S2)}$$

$$\partial_t V = \nabla\cdot\big(D_V V\nabla V\big) + p_V \frac{k_{\mathrm{PV}}}{k_{\mathrm{PV}}+F} I - c_V V, \quad \textbf{(S3)}$$

$$\partial_t F = \nabla\cdot\big(D_F F\nabla F\big) + p_F I\Big(1+\frac{a_F F^2}{K_F^2+F^2}\Big) - c_F F. \quad \textbf{(S4)}$$

Zero-flux (Neumann) boundary conditions are imposed on $\partial\Omega$. Susceptible cells become infected at a baseline rate $\beta VS$, and infected cells die at rate $\delta_I$. Infected cells produce virions and IFN at rates $p_V$ and $p_F$, respectively, while extracellular virions and IFN are cleared at rates $c_V$ and $c_F$. Four IFN-dependent mechanisms are considered:

- *Blocking of infection* (BI): the factor $k_{\mathrm{IF}}/(k_{\mathrm{IF}}+F)$ reversibly reduces the infection rate, with half-maximal inhibition at $F = k_{\mathrm{IF}}$;
- *Refractory conversion* (RF): the term $-k_R F/(K_R+F)S$ irreversibly removes susceptible cells from the infection process, with maximum conversion rate $k_R$ and half-maximal concentration $K_R$;
- *Blocking of virion production* (BP): the factor $k_{\mathrm{PV}}/(k_{\mathrm{PV}}+F)$ reduces virion production by infected cells, with half-maximal inhibition at $F = k_{\mathrm{PV}}$;
- *Positive feedback* (PF): the factor $1+a_F F^2/(K_F^2+F^2)$ enhances IFN production by up to $(1+a_F)$-fold, with half-maximal enhancement at $F = K_F$.

Virions and IFN spread through porous-medium diffusion rather than linear diffusion:

$$\nabla\cdot(D_X X^{m_X}\nabla X) = \frac{D_X}{m_X+1}\Delta X^{m_X+1}, \quad \textbf{(S5)}$$

with $m_V = m_F = 1$, giving $\frac{1}{2}D_X\Delta(X^2)$. Unlike linear diffusion, porous-medium diffusion permits finite-speed propagation from initially localized, compactly supported distributions, avoiding instantaneous spreading across the domain. We assume $D_F = 100 D_V$, motivated by the smaller hydrodynamic size of IFN molecules (a few nm) relative to influenza virions ($\sim$100 nm). This ratio is an approximation based on inverse size scaling; the absolute diffusivities depend on the unspecified physical length of the normalized domain. Numerically, the diffusion operator is evaluated as $\frac{1}{2}D_X\Delta\big(\max(X,0)+\varepsilon\big)^2$, with $\varepsilon = 10^{-6}$ to regularize the operator near zero concentrations.

Initially, susceptible cells are uniformly distributed, $S(\mathbf{x},0) = 100$, with $I(\mathbf{x},0) = F(\mathbf{x},0) = 0$. Infection is initiated by a Gaussian virion distribution,

$$V(\mathbf{x},0) = \exp\left(-\frac{\|\mathbf{x}-\mathbf{x}_c\|^2}{2\ell^2}\right), \tag{S6}$$

centered at $\mathbf{x}_c = (\frac{1}{2}, \frac{1}{2})$ with $\ell = 0.04$.

**Model variants and the uninfected fraction.** The 16 possible combinations of IFN mechanisms are generated by selectively disabling terms in the full model. BI and BP are disabled by setting $k_{\mathrm{IF}} \to \infty$ and $k_{\mathrm{PV}} \to \infty$, respectively, so their inhibition factors approach unity. RF is disabled by setting $k_R = 0$, and PF by setting $a_F = 0$. Numerical implementations use sentinel values of $10^{12}$ and $10^{-12}$, respectively.
The PF-only combination is excluded because, in the absence of BI, BP, and RF, IFN does not influence the dynamics of $S$, $I$, or $V$. Its viral trajectory is therefore identical to that of the null model, differing only in its IFN predictions and corresponding likelihood contribution. The remaining 15 combinations are listed in Table S2.
Refractory cells are not explicitly tracked because their abundance can be recovered from cell conservation. Defining the spatial integral as $\langle g\rangle(t) = \int_\Omega g(\mathbf{x},t)\,\mathrm{d}\mathbf{x}$, summing the susceptible and infected cell equations gives

$$\frac{\mathrm{d}}{\mathrm{d}t}\langle S+I\rangle = -\left\langle k_R \frac{F}{K_R+F} S\right\rangle - \delta_I\langle I\rangle. \tag{S7}$$

The fraction of cells that have never been infected, comprising susceptible and refractory cells, is therefore

$$f_u(t) = 1 - \frac{\langle I\rangle(t) + \int_0^t \delta_I \langle I\rangle(s)\,\mathrm{d}s}{\langle S\rangle(0)}, \tag{S8}$$

where the integral accounts for cumulative infection-induced cell death. During constrained calibration, parameter sets yielding $f_u(10) < 0.8$ are rejected.

**Calibration and numerics.** The infected-cell death rate, $\delta_I = 1.8$ d$^{-1}$, and virion clearance rate, $c_V = 4.2$ d$^{-1}$, were fixed from ref. (1). The IFN diffusivity $D_F$ was set relative to $D_V$, leaving 7–11 free parameters depending on the IFN mechanisms included (Table S3). Model observables were the spatial integrals $g_V(t) = \langle V\rangle(t)$ and $g_F(t) = \langle F\rangle(t)$, fitted to experimental data using the same censored-Gaussian negative log-likelihood and profiled offsets $s_V$ and $s_F$ as in the agent-based model (section 2).
The equations were discretized on a uniform $16 \times 16$ grid using a five-point Laplacian with Neumann boundary conditions and integrated with a sparse-Jacobian backward differentiation formula (BDF) solver (relative tolerance $10^{-4}$; absolute tolerance $10^{-6}$). Parameters were optimized by differential evolution in $\log_{10}$ space, using a population of 15 per parameter, up to 500 generations, a convergence tolerance of 0.005, deferred updating, and no final polishing.

**Results.** All 15 IFN mechanism combinations reproduced the viral and IFN kinetics with nearly identical likelihoods. Negative log-likelihoods differed by only 0.018 units, compared with a variation of 0.09–0.23 across optimization seeds (Fig. S1, Table S2). However, the fraction of cells spared from infection by day 10 varied from 0% to 64% among these fits, indicating that the observed kinetics did not uniquely constrain tissue damage.
Imposing $f_u(10) \geq 0.8$ favored refractory conversion alone (M05), while the null model was penalized by more than 100 negative log-likelihood units. Nevertheless, every constrained model attained the same likelihood as its unconstrained counterpart. Spatial distributions (Fig. S1*F*) showed that tissue sparing resulted from widespread refractory conversion or low infection throughout the domain, rather than spatial confinement of infection. In all models, the infection front reached the entire domain by day 3.
When estimated freely, the IFN clearance rate converged to $c_F \approx 0.3$ d$^{-1}$, corresponding to a half-life of approximately 2.3 d. Fixing $c_F = 2.0$ d$^{-1}$ increased the negative log-likelihood by 15.9 units. This slow effective clearance may reflect processes not explicitly represented in the model, such as distinct IFN transcript and protein dynamics or sustained IFN production by recruited immune cells, rather than the intrinsic decay rate of secreted IFN-$\beta$.

#### 2. The agent-based model

**Tissue network and field update.** The epithelial tissue is represented as a network in which each cell $i$ is connected to all cells $j$ within a distance $d_{ij} \leq r$, where $r$ is the neighborhood radius and $d_{ij}$ is the distance between cells $i$ and $j$. Connections are weighted by $w_{ij} = 1/d_{ij}^2$. At each time step ($\Delta t = 1$ min), the extracellular fields $X \in \{V, F\}$ are updated according to

$$X_i \leftarrow (1-\delta_X)\left[X_i + D_X \sum_j w_{ij}(X_j - X_i)\right], \tag{S9}$$

where $\delta_X$ is the per-step clearance fraction and $D_X$ controls transport between neighboring cells. The update is stable and preserves nonnegative concentrations when $D_X \max_i \sum_j w_{ij} \leq 1$.

**Cell states.** Initially, each cell is assigned a non-permissive state with probability $p_{\mathrm{np}}$ or is otherwise susceptible. Susceptible cells can become refractory or infected, while infected cells eventually die. Infected cells are classified as IFN-competent ($\mathrm{I}^+$) or IFN-incompetent ($\mathrm{I}^-$).

Cell-state transitions are evaluated at each time step using Hill functions,

$$H(x;K) = \frac{x^n}{K^n + x^n}. \tag{S10}$$

A susceptible cell becomes refractory with probability $H(\log_{10}(F_i+1);K_R)$ (RF). Otherwise, it becomes infected with probability

$$H(\log_{10}(V_i+1);K_V)\left[1 - H(\log_{10}(F_i+1);K_{BI})\right], \tag{S11}$$

where the bracket represents IFN-mediated blocking of infection (BI). Newly infected cells enter the $\mathrm{I}^+$ state with probability $p_+$ and the $\mathrm{I}^-$ state otherwise. Infected-cell lifetimes are sampled from a Poisson distribution with mean $\tau_I$ converted from minutes to hours. The sampled lifetimes are then converted back to minutes for the simulation.

**Virion and IFN production.** Virion production depends on the time since infection, $a$, and local IFN concentration:

$$\dot{V}_i(a) = \left[1 - H(F_i;K_{BP})\right] \frac{\mathrm{d}V_{\mathrm{cum}}}{\mathrm{d}a}, \tag{S12}$$

where the cumulative single-cell virion production follows

$$V_{\mathrm{cum}}(a) = \frac{A\,a^{m_p}}{K_p^{m_p} + a^{m_p}}. \tag{S13}$$

This function was fitted to the single-cell production data of ref. (2), yielding $A = 17{,}704$ virions, $K_p = 16.1$ h, and $m_p = 2.39$. The inhibitory factor represents blocking of virion production (BP).

Both $\mathrm{I}^+$ and $\mathrm{I}^-$ cells release virions into their local extracellular field, but only $\mathrm{I}^+$ cells produce IFN. IFN production is mediated by an intracellular dsRNA variable, $\psi_i$, updated as

$$\psi_i \leftarrow \psi_i + k_{\mathrm{syn}}\dot{V}_i - k_{\mathrm{deg}}\psi_i, \tag{S14}$$

$$F_i \leftarrow F_i + p_F \psi_i \left[1 + \alpha_{PF} H(F_i;K_{PF})\right], \tag{S15}$$

where the bracket represents IFN-mediated positive feedback (PF). Unlike RF and BI, BP and PF depend on the untransformed IFN concentration. RF and BI regulate susceptible-cell transitions, where local IFN levels span several orders of magnitude across the tissue. Using untransformed IFN would therefore produce an almost step-like response near the lesion boundary. In contrast, BP and PF act within infected cells, where IFN levels vary over a narrower range ($\sim$1.5 orders of magnitude), making a half-maximal response based on untransformed IFN more appropriate.

**Cytotoxic T lymphocytes.** The adaptive immune response is represented by a single systemic cytotoxic T lymphocyte (CTL) pool, $T$, initialized at $T_0$. At each time step, the pool is updated according to

$$T \leftarrow T + \rho_T H(\bar{F};K_{\bar{F}})T\left(1 - \frac{T}{T_{\mathrm{max}}}\right) - \delta_T T, \tag{S16}$$

where $\bar{F}$ is the mean extracellular IFN concentration across all cells. Each infected cell is killed with probability $H(T;K_T)$ per time step. At each time step, CTL-mediated killing is evaluated before natural infected-cell death.

**Initial conditions, observables, and implementation.** Initially, all cells have zero IFN and intracellular dsRNA. Infection is initiated by depositing $V_0 = 100$ virions on each of $N_0$ distinct, randomly selected cells. For the two-dimensional section z0, $N_0 = 38$, corresponding to the number of IAV-positive clusters identified by HDBSCAN (Fig. S2). For the three-dimensional tissue, $N_0 = 100$.

Model observables are the total extracellular virion and IFN levels, $g_V(t) = \sum_i V_i(t)$ and $g_F(t) = \sum_i F_i(t)$, averaged across simulation replicates. Lesion extent is defined as the fraction of all cells that are infected or dead at day 2.

TissueFlu is implemented in CUDA C/C++, with one GPU thread per cell. Simulations are reproducible from their random seeds. Fixed and calibrated parameters are listed in Table S5.

**Calibration.** Model predictions were compared with experimental data using a censored-Gaussian negative log-likelihood with profiled log-scale offsets $s_V$ and $s_F$:

$$J_V = \frac{1}{2\sigma_V^2}\sum_d \sum_{k\in\det(d)} \left(\log_{10} V_{d,k} - \log_{10} g_V(d) - s_V\right)^2 - \sum_d \sum_{k\in\mathrm{cen}(d)} \log\Phi\left(\frac{\log_{10} L - \log_{10} g_V(d) - s_V}{\sigma_V}\right), \tag{S17}$$

$$J_F = \frac{1}{2\sigma_F^2}\sum_{d\in\{2,4,6\}} \sum_k \left(\log_{10} F_{d,k} - \log_{10} g_F(d) - s_F\right)^2, \tag{S18}$$

where $V_{d,k}$ and $F_{d,k}$ are experimental measurements at day $d$, $\det(d)$ and $\mathrm{cen}(d)$ denote detected and left-censored viral observations, respectively, $L$ is the viral detection limit, and $\Phi$ is the standard normal cumulative distribution function. The residual standard deviations were fixed at $\sigma_V = 0.62$ and $\sigma_F = 0.16$. The objective was minimized over the offsets, $J = \min_{s_V,s_F}(J_V + J_F)$.

Models were ranked using the corrected Akaike information criterion (AICc), with $n = 50$ observations and $k$ equal to the number of calibrated parameters plus the two offsets. Calibration proceeded in stages: Latin-hypercube sampling of the free parameter space, local refinement within trust regions around the best candidates (guided by a Gaussian-process surrogate in the initial campaigns), and confirmation of the final parameter set using 30–50 simulation replicates (Fig. S3).

The neighborhood radius was screened over $r \in \{4,5,6,8,10,12,14,16\}$ using the day-2 lesion extent, yielding $r = 10$ for the two-dimensional section and $r = 6$ for the three-dimensional tissue.

**The search space.** Table S5 reports the parameter search bounds and the values selected by each calibration. All free parameters were sampled log-uniformly because they span several orders of magnitude and their characteristic scales were not known a priori. Three aspects of the search design warrant clarification.

First, the transport coefficients $D_V$ and $D_F$ were not estimated independently. Instead, the ratio $D_F/D_V$ was sampled within $[1,10]$, and $D_F$ was calculated as $\min\{(D_F/D_V)D_V, 0.95D_{\max}\}$. The upper cap ensures numerical stability of the explicit field update (Eq. S9), which requires $D_X \max_i \sum_j w_{ij} \le 1$. This gives $D_{\max} = 0.096$ for $r = 10$ and 0.047 for $r = 6$.

Unlike the continuum model, where $D_F = 100D_V$ was assumed based on molecular size, $D_X$ in the ABM is a discrete transport coefficient rather than a physical diffusivity. Both fields propagate through the same network, using identical neighborhood radii and edge weights, so the ratio controls their relative rates of spatial redistribution rather than their connectivity. We therefore allowed the data to determine this ratio instead of imposing the continuum-model assumption. The fitted ratios were 1.6 and 5.0 for the two-dimensional section (with and without CTL, respectively) and 5.8, 1.3, and 2.1 for the three-dimensional tissue. None reached the search bounds, although $D_F$ reached the numerical stability cap in the two-dimensional fit without CTL.

Second, three parameters were fixed to reduce the dimensionality of the calibration. The infection threshold $K_V$ was fixed after a parameter sweep showed negligible sensitivity of the model outcomes to its variation. Its value was selected to reproduce the experimental day-2 lesion extent in the null model, without IFN-mediated protection or CTL activity. The mean infected-cell lifetime $\tau_I$ and virion clearance $\delta_V$ were fixed using values informed by refs. (2) and (1), respectively. Fixing these parameters also reduced potential confounding with transport parameters when fitting bulk viral kinetics. The exception was the three-dimensional calibration without CTL, for which no satisfactory fit was obtained with $\tau_I$ and $\delta_V$ fixed. Allowing both parameters to vary improved the fit, and the additional free parameters were included in the AICc penalty.

Third, $k_{\mathrm{syn}}$ and $p_F$ were fixed to 1 to reduce parameter redundancy. Because IFN measurements are reported as fold changes, the absolute scale of the modeled IFN field is not directly constrained by the data and is accommodated by the profiled offset $s_F$. Fixing these parameters establishes a reference scale for IFN-related thresholds $K_R$, $K_{BI}$, $K_{BP}$, $K_{PF}$, and $K_{\bar{F}}$.

#### 3. Network characterization and the homogeneous control

For each tissue structure and neighborhood radius, we constructed the weighted network used in the simulations and characterized its degree and strength ($s_i = \sum_j w_{ij}$) distributions, connected components, and mean local clustering coefficient (3,000 sampled cells). We also measured the number of cells $B(k)$ reachable within $k$ network steps from randomly selected starting cells in the largest connected component (50 cells per structure, or 10 for the three-dimensional tissue). The effective network dimension was estimated from the slope of $\log B(k)$ versus $\log k$ over $3 \le k \le 20$ (Table S6).

Tissue holes were quantified using cubical persistent homology implemented in GUDHI (3). We computed the distance from each point on a unit grid to the nearest cell and analyzed its sublevel sets. At threshold $r/2$, the sublevel set corresponds to the union of balls of radius $r/2$ centered on the cells, with overlaps determined by the neighborhood radius $r$. Hole size was defined by the radius at which a hole disappeared, and short-lived features persisting less than 1.5 times their appearance radius were excluded to remove small gaps between adjacent balls. Across tissue sections, hole density peaked at 335–372 per $10^4$ cells for $r = 9$–$12$. Because the spacing between stacked sections was assumed to be one spatial unit, we assessed sensitivity to this assumption by rescaling the $z$ coordinate by factors of 2, 3, and 5 and recomputing network and hole measures in $1200 \times 1200$ and $400 \times 400$ windows spanning all sections (Table S8).

The homogeneous control consisted of a hexagonal lattice with 233,479 cells, matching section z0. The lattice spacing was set to $a = 2.57$ and the neighborhood radius to $r = 6$, giving each interior cell 18 neighbors at distances $a$, $\sqrt{3}a$, and $2a$. The spacing was chosen to match the mean network strength of section z0 at $r = 10$.

#### 4. Results of the agent-based model

This section presents additional agent-based model results supporting the main text. Results already reported in the main text are repeated only when needed for comparison.

**Model selection on section z0.** All 15 combinations of the four IFN mechanisms were calibrated under three scenarios, with model rankings summarized in Table S4. When calibrated to days 0–6, covering viral growth and peak infection, the mechanism-free model provided the best feasible fit ($J = 43.5$). Although refractory conversion alone achieved a slightly lower cost ($J = 41.8$), it exceeded the experimental day-2 lesion extent (18.3% versus $13 \pm 5\%$) and was therefore excluded. All other combinations had $\Delta\text{AICc} \geq 4$ relative to the mechanism-free model. Thus, early viral and IFN kinetics did not support the inclusion of additional IFN mechanisms.

Over the full ten-day period without CTLs, the combination BI+RF+BP was favored ($J = 89.7$), with the mechanism-free model 26.7 AICc units behind. However, even the best-fitting combination reproduced the data substantially less well than models including CTLs. With CTLs, the mechanism-free model was again preferred ($J = 44.5$), and all additional IFN mechanisms were penalized by $\Delta\text{AICc} \geq 5$.

Together, these results indicate that IFN-mediated mechanisms are favored primarily when the model must reproduce viral decline without adaptive immunity, but they remain insufficient to account for the observed clearance dynamics.

**IFN mechanisms alone and in pairs.** Figure S4 shows the day-2 lesion extent from the single-mechanism sweep whose day-10 tissue damage is presented in Fig. 4*A* of the main text. Damage decreases progressively with increasing mechanism strength, but only refractory conversion shifts the day-2 lesion outside the experimental range, with an abrupt change between consecutive strength levels. Figure S5 extends this analysis to pairwise combinations at three strength levels per mechanism. Positive feedback enhances the protective effect of refractory conversion, reducing tissue damage beyond that achieved by either mechanism alone, but provides little additional protection when combined with blocking of infection.

**Interferon without CTLs, on real tissue and on the lattice.** Figure S6 compares 400 parameter sets sampled across IFN and transport parameters, with all four IFN mechanisms active and CTLs absent. Each parameter set was simulated on both section z0 and the matched homogeneous lattice. Real tissue preserved more cells than the lattice in 395 of 400 cases. On section z0, 299 parameter sets spared at least 80% of permissive cells, but these cases largely represented failure to establish infection rather than spatial containment. Their median day-2 lesion extent was only 0.14% (maximum 4.4%), well below the experimental $13 \pm 5\%$.

On the lattice, 243 parameter sets spared at least 80% of permissive cells, but only five also reproduced the measured day-2 lesion extent. In these cases, protection was associated with widespread refractory conversion, with a median of 44% of cells refractory by day 10. Thus, homogeneous tissue could reproduce both the early lesion extent and substantial tissue sparing, but only through widespread IFN-mediated protection that was not supported by the IFN measurements.

To isolate the effect of tissue architecture, each calibrated parameter set from section z0 was re-simulated on the lattice without modification (Table S7). Movies S1 and S2 illustrate the resulting spatial dynamics: infection foci expand radially and merge on the lattice, whereas airspaces restrict their expansion and maintain spatial separation on the tissue section.

**Robustness of the calibrated section model.** We assessed the sensitivity of the calibrated z0 model to neighborhood radius and tissue structure. Figure S7 compares simulations across neighborhood radii, with diffusivities either held fixed or rescaled to preserve the mean coupling strength. The rescaled results indicate that infection outcomes depend primarily on the number of connected neighbors rather than coupling strength. Figure S8 applies the calibrated z0 parameters unchanged to three additional sections of the same lung. Viral trajectories show similar temporal patterns, with differences reflecting section-specific connectivity. The least connected section (z15; mean degree 12.3 versus 17.0 in z0) exhibits the lowest cell loss.

### 5. Supplementary tables

**Table S1.** Spatial models of acute viral infections of epithelial tissue. All but one address respiratory viruses; the herpes simplex virus-2 genital-ulcer model (4) is listed for its agent-based treatment of tissue-resident lymphocytes. Method: partial differential equations (PDE), agent-based models (ABM), cellular automata (CA), hybrid PDE–ABM, or an age-dependent branching process. Domain: the geometry on which the model is solved; "space-free" denotes a model that represents two spatial scales without spatial coordinates. Calibration: fitted to bulk or spatially aggregated measurements (nonspatial), to spatially resolved measurements (spatial), or fixed from literature without fitting (fixed).

| **Reference** | **Virus** | **Domain** | **Method** | **Calibration** |
|---|---|---|---|---|
| (5) | SARS-CoV-2 | Airway surrogate | ABM/CA | Fixed |
| (6) | SARS-CoV-2 | Airway surrogate | PDE | Nonspatial |
| (7) | SARS-CoV-2 | Tissue culture surrogate | Hybrid | Nonspatial |
| (8) | SARS-CoV | Tissue culture surrogate | ABM/CA | Spatial, Nonspatial |
| (9) | Generic | Tissue culture surrogate | PDE | Fixed |
| (10) | Influenza A | Tissue culture surrogate | ABM/CA | Fixed |
| (11) | Influenza A | Tissue culture surrogate | ABM/CA | Fixed |
| (12) | SARS-CoV-2; IAV | Tissue culture surrogate | Hybrid | Fixed |
| (13) | SARS-CoV-2 | Tissue culture surrogate | PDE | Fixed |
| (14) | SARS-CoV-2 | Tissue culture surrogate | PDE | Fixed |
| (15) | Influenza A | Airway surrogate | PDE | Fixed |
| (16) | SARS-CoV-2 | Airway surrogate | ABM/CA | Fixed |
| (17) | Influenza A | Tissue culture surrogate | ABM/CA | Fixed |
| (18) | Influenza A | Tissue culture surrogate | Hybrid | Nonspatial, Fixed |
| (19) | Generic | Tissue culture surrogate | Hybrid | Fixed |
| (20) | Influenza A | Tissue culture surrogate | Hybrid | Fixed |
| (21) | Influenza A | Tissue culture surrogate | ABM/CA | Nonspatial |
| (22) | SARS-CoV-2; IAV | Space-free (foci + long-range seeding) | Branching process; CA | Fixed |
| (23) | HSV-1 | Tissue culture surrogate | ABM/CA | Nonspatial |
| (4) | HSV-2 | Genital epithelium surrogate (2-D lattice) | ABM/CA | Nonspatial |
| This work | Influenza A | Histology-derived lung (2-D, 3-D) | PDE; ABM | Nonspatial + spatial |

**Table S2.** Reaction–diffusion models calibrated without and with the sparing constraint $f_u(10) \geq 0.8$. BI, blocking of infection; BP, blocking of virion production; RF, refractory conversion; PF, positive feedback. $k$ counts calibrated parameters plus the two offsets; $n = 50$. NLL, negative log-likelihood; ΔAICc within each calibration. The NLL spread between optimizer seeds is 0.09–0.23.

| | | | Unconstrained | | | $f_u(10) \geq 0.8$ | | |
|---|---|---|---|---|---|---|---|---|
| Model | Mechanisms | $k$ | NLL | ΔAICc | $f_u(10)$ | NLL | ΔAICc | $f_u(10)$ |
| M01 | – | 9 | 37.287 | 0.00 | 0.000 | 153.152 | 228.55 | 0.800 |
| M05 | RF | 10 | 37.288 | 3.14 | 0.001 | 37.307 | 0.00 | 0.802 |
| M02 | BI | 10 | 37.291 | 3.15 | 0.000 | 52.319 | 30.02 | 0.800 |
| M03 | BP | 10 | 37.295 | 3.16 | 0.000 | 47.643 | 20.67 | 0.800 |
| M06 | BI+RF | 11 | 37.287 | 6.45 | 0.005 | 37.309 | 3.31 | 0.814 |
| M10 | BI+PF | 11 | 37.287 | 6.45 | 0.000 | 37.359 | 3.41 | 0.856 |
| M11 | BP+PF | 11 | 37.289 | 6.45 | 0.000 | 50.113 | 28.92 | 0.802 |
| M04 | BI+BP | 11 | 37.290 | 6.45 | 0.000 | 41.886 | 12.46 | 0.803 |
| M07 | BP+RF | 11 | 37.291 | 6.46 | 0.000 | 37.311 | 3.31 | 0.835 |
| M13 | RF+PF | 11 | 37.292 | 6.46 | 0.033 | 37.315 | 3.32 | 0.809 |
| M14 | BI+RF+PF | 12 | 37.290 | 9.94 | 0.636 | 37.307 | 6.79 | 0.824 |
| M08 | BI+BP+RF | 12 | 37.293 | 9.94 | 0.272 | 37.305 | 6.79 | 0.802 |
| M15 | BP+RF+PF | 12 | 37.299 | 9.96 | 0.055 | 37.311 | 6.80 | 0.809 |
| M12 | BI+BP+PF | 12 | 37.305 | 9.97 | 0.000 | 37.481 | 7.14 | 0.816 |
| M16 | BI+BP+RF+PF | 13 | 37.305 | 13.65 | 0.001 | 37.306 | 10.47 | 0.812 |

**Table S3.** Reaction–diffusion model: search bounds and estimates. Units: days and the unit square; $S(0) = 100$. $^\dagger$fixed or derived ($D_F = 100D_V$); "off", mechanism disabled.

| | Meaning | Bounds | M05, constr. | M01, unconstr. | M05, unconstr. |
|---|---|---|---|---|---|
| $\beta$ | infection rate | $8.00 \times 10^{-5}$–8.00 | $2.06 \times 10^{-4}$ | $1.14 \times 10^{-4}$ | $1.13 \times 10^{-4}$ |
| $p_V$ | virus production rate | 1.00–$1.00 \times 10^{4}$ | 9020.38 | $1.49 \times 10^{4}$ | $1.55 \times 10^{4}$ |
| $D_V$ | virus diffusion coefficient | $5.00 \times 10^{-10}$–$5.00 \times 10^{-3}$ | $1.32 \times 10^{-6}$ | $1.69 \times 10^{-7}$ | $1.48 \times 10^{-7}$ |
| $p_F$ | IFN production rate | $3.00 \times 10^{-3}$–100.00 | 0.12 | 0.58 | 0.06 |
| $c_F$ | IFN clearance rate | $1.00 \times 10^{-3}$–20.00 | 0.30 | 0.31 | 0.31 |
| $k_{\text{IF}}$ | IFN half-max, blocks-infection (BI) | $2.00 \times 10^{-3}$–20.00 | off | off | off |
| $k_{\text{PV}}$ | IFN half-max, blocks-production (BP) | 0.10–1000.00 | off | off | off |
| $k_R$ | refractory conversion rate | 0.05–500.00 | 14.84 | off | 1.09 |
| $K_R$ | IFN half-max for RF | 0.01–100.00 | 0.07 | 15.95 | 45.59 |
| $a_F$ | positive-feedback fold-enhancement | 10.00–$1.00 \times 10^{5}$ | off | off | off |
| $K_F$ | IFN half-max for PF | 0.01–100.00 | 1.58 | 3.04 | 7.20 |
| $\delta_I$ | infected-cell death rate | 0.30–30.00 | $1.80^\dagger$ | $1.80^\dagger$ | $1.80^\dagger$ |
| $c_V$ | virus clearance rate | 0.10–10.00 | $4.20^\dagger$ | $4.20^\dagger$ | $4.20^\dagger$ |
| $D_F$ | IFN diffusion coefficient | – | $1.32 \times 10^{-4\dagger}$ | $1.69 \times 10^{-5\dagger}$ | $1.48 \times 10^{-5\dagger}$ |

**Table S4.** Agent-based model selection on section z0. Each of the 15 combinations of the four IFN mechanisms was calibrated in each scenario; $J$ is the fit cost on days 0–10, $f_2$ the day-2 lesion extent (target $13.0 \pm 5.0\%$), feasible: day-2 lesion (and T-cell windows when CTL is on) satisfied. $\Delta$AICc relative to the best feasible model.

| Scenario | Model | $k$ | $J$ | $f_2$ (%) | Feasible | $\Delta$AICc |
|---|---|---|---|---|---|---|
| no CTL, days 0–6 | RF | 7 | 41.78 | 18.3 | no | -0.46 |
| | none | 6 | 43.48 | 17.3 | yes | 0.00 |
| | BI | 7 | 44.07 | 17.2 | yes | 4.12 |
| | BP | 7 | 44.65 | 17.4 | yes | 5.29 |
| | RF+BP | 8 | 44.61 | 15.6 | yes | 8.35 |
| | BP+PF | 8 | 44.65 | 17.8 | yes | 8.43 |
| | BI+BP | 8 | 44.97 | 17.1 | yes | 9.07 |
| | BI+PF | 8 | 45.22 | 16.3 | yes | 9.57 |
| | BI+RF+BP | 9 | 43.57 | 15.3 | yes | 9.63 |
| | BI+BP+PF | 9 | 43.80 | 17.9 | yes | 10.09 |
| | RF+PF | 8 | 46.15 | 17.4 | yes | 11.43 |
| | BI+RF+PF | 9 | 45.52 | 12.7 | yes | 13.53 |
| | BI+RF | 8 | 48.29 | 12.2 | yes | 15.72 |
| | BI+RF+BP+PF | 10 | 50.73 | 11.4 | yes | 27.53 |
| | RF+BP+PF | 9 | 53.38 | 12.9 | yes | 29.25 |
| no CTL, days 0–10 | BI+RF+BP | 9 | 89.74 | 16.4 | yes | 0.00 |
| | BI+RF+PF | 9 | 91.74 | 16.4 | yes | 4.00 |
| | BI | 7 | 101.04 | 19.2 | no | 16.78 |
| | RF+PF | 8 | 100.03 | 17.0 | yes | 17.60 |
| | BI+BP | 8 | 103.74 | 18.0 | no | 25.02 |
| | BP | 7 | 105.42 | 17.9 | yes | 25.54 |
| | BI+BP+PF | 9 | 102.67 | 19.2 | no | 25.87 |
| | RF | 7 | 105.59 | 15.0 | yes | 25.87 |
| | none | 6 | 107.35 | 18.4 | no | 26.68 |
| | BP+PF | 8 | 105.43 | 18.0 | no | 28.39 |
| | BI+PF | 8 | 108.01 | 17.0 | yes | 33.56 |
| | RF+BP+PF | 9 | 111.82 | 16.2 | yes | 44.17 |
| | RF+BP | 8 | 114.65 | 15.8 | yes | 46.85 |
| | BI+RF | 8 | 123.70 | 12.4 | yes | 64.93 |
| | BI+RF+BP+PF | 10 | 122.95 | 16.7 | yes | 69.57 |
| CTL, days 0–10 | none | 9 | 44.53 | 8.4 | yes | 0.00 |
| | BI | 10 | 45.56 | 9.2 | yes | 5.19 |
| | BP | 10 | 47.73 | 16.7 | yes | 9.53 |
| | RF+BP+PF | 12 | 45.86 | 18.0 | no | 12.59 |
| | BI+RF+PF | 12 | 47.30 | 16.0 | yes | 15.47 |
| | BI+BP+PF | 12 | 47.33 | 12.3 | yes | 15.53 |
| | BP+PF | 11 | 52.15 | 8.6 | yes | 21.69 |
| | BI+PF | 11 | 58.34 | 15.6 | yes | 34.06 |
| | RF+PF | 11 | 58.38 | 15.0 | yes | 34.13 |
| | BI+BP | 11 | 58.49 | 15.8 | yes | 34.37 |
| | BI+RF+BP+PF | 13 | 55.24 | 11.8 | yes | 35.02 |
| | RF+BP | 11 | 65.07 | 14.8 | yes | 47.52 |
| | BI+RF+BP | 12 | 63.46 | 15.0 | yes | 47.79 |
| | BI+RF | 11 | 70.81 | 11.1 | yes | 59.00 |
| | RF | 10 | 75.15 | 9.9 | yes | 64.38 |

**Table S5.** Agent-based model: search ranges and the calibrated points used in the main text. Ranges are the bounds of the design; every searched parameter was sampled log-uniformly between them, and "fixed" marks a parameter that was not part of the design. Fixed in all calibrations: $n = 2$ for every Hill function, $T(0) = 10^{-4}$, $T_{\max} = 1000$; virion release kinetics from ref. (2). One time step is one minute; distances in cell diameters. $k_{\text{syn}}$ and $p_F$ were fixed to 1 and define the arbitrary IFN unit of the model. §chosen before calibration from a screen of the day-2 lesion extent over $r \in \{4, 5, 6, 8, 10, 12, 14, 16\}$. ¶the number of IAV-positive clusters of the tissue (Fig. S2). ‖low end extended to $10^{-4}$ (3-D, no CTL) and $10^{-5}$ (3-D, CTL) for $D_V$, and to $10^{-4}$ for $\delta_F$ in both, because without CTL, or without the IFN mechanisms, the day-2 lesion can only be held in its measured band by slower transport. ** $D_F$ is not searched independently: the free parameter is the ratio $D_F/D_V \in [1, 10]$, and $D_F$ is then capped at $0.95$ times the stability limit of Eq. [S9] ($0.096$ at $r = 10$, $0.047$ at $r = 6$). ††fixed at 1800 and $2.34 \times 10^{-3}$ everywhere except the 3-D calibration without CTL, which had no feasible region until these two were also freed.

| | Meaning | Search range | z0, CTL | z0, no CTL | 3-D, CTL | 3-D, no CTL |
|---|---|---|---|---|---|---|
| $r$ | neighbourhood radius (cell diameters) | screened§ | 10 | 10 | 6 | 6 |
| $N_0$ | inoculated cells | fixed¶ | 38 | 38 | 100 | 100 |
| | virions per inoculated cell | fixed | 100 | 100 | 100 | 100 |
| $p_{\text{NP}}$ | non-permissive fraction | fixed | 0.200 | 0.200 | 0.200 | 0.200 |
| | IFN-competent fraction of infected cells | fixed | 0.500 | 0.500 | 0.500 | 0.500 |
| $K_V$ | infection half-max ($\log_{10}$ virions) | fixed | 3.50 | 3.50 | 3.50 | 3.50 |
| $\tau_I$ | mean infected-cell lifetime (min) | 600–4320†† | 1800 | 1800 | 1800 | 3210 |
| $\delta_V$ | virion clearance ($\text{min}^{-1}$) | $5 \times 10^{-4}$–$10^{-2}$†† | $2.34 \times 10^{-3}$ | $2.34 \times 10^{-3}$ | $2.34 \times 10^{-3}$ | $2.64 \times 10^{-3}$ |
| $D_V$ | virion diffusivity | $5 \times 10^{-4}$–$2 \times 10^{-2}$‖ | $1.26 \times 10^{-3}$ | 0.018 | $1.39 \times 10^{-4}$ | 0.020 |
| $D_F$ | IFN diffusivity | $(1–10)\, D_V$** | $1.97 \times 10^{-3}$ | 0.091 | $8.03 \times 10^{-4}$ | 0.026 |
| $\delta_F$ | IFN clearance ($\text{min}^{-1}$) | $2 \times 10^{-4}$–$5 \times 10^{-3}$‖ | $3.49 \times 10^{-4}$ | $2.81 \times 10^{-4}$ | $1.72 \times 10^{-4}$ | $4.21 \times 10^{-4}$ |
| $k_{\text{syn}}$ | dsRNA synthesis (per virion released) | fixed | 1 | 1 | 1 | 1 |
| $k_{\text{deg}}$ | dsRNA degradation ($\text{min}^{-1}$) | $10^{-4}$–$10^{-1}$ | $1.18 \times 10^{-4}$ | $1.28 \times 10^{-3}$ | 0.015 | 0.063 |
| $p_F$ | IFN production (per dsRNA per min) | fixed | 1 | 1 | 1 | 1 |
| BI | blocking of infection on/off | fixed | off | on | off | on |
| $K_{BI}$ | blocking-of-infection half-max ($\log_{10}$ IFN) | 1–100 | – | 76.34 | – | 47.12 |
| RF | refractory conversion on/off | fixed | off | on | off | on |
| $K_R$ | refractory half-max ($\log_{10}$ IFN) | 1–100 | – | 53.32 | – | 4.79 |
| BP | blocking of production on/off | fixed | off | on | off | on |
| $K_{BP}$ | BP half-max (IFN) | $10^2$–$10^8$ | – | $6.14 \times 10^6$ | – | 2045.79 |
| PF | positive feedback on/off | fixed | off | off | off | on |
| $K_{PF}$ | PF half-max (IFN) | $10^2$–$10^8$ | – | – | – | $2.08 \times 10^6$ |
| $\alpha_{PF}$ | PF amplification | fixed | – | – | – | 1 |
| CTL | CTL compartment on/off | fixed | on | off | on | off |
| $\rho_T$ | CTL expansion ($\text{min}^{-1}$) | $5 \times 10^{-4}$–$10^{-2}$ | $1.03 \times 10^{-3}$ | – | $1.61 \times 10^{-3}$ | – |
| $K_{\text{IFN}}$ | CTL expansion half-max (mean IFN) | $10^{-2}$–$10^6$ | 0.105 | – | 1597.65 | – |
| $K_T$ | CTL killing half-max | 0.1–500 | 4.44 | – | 9.82 | – |
| $\delta_T$ | CTL decay ($\text{min}^{-1}$) | fixed | $7.64 \times 10^{-6}$ | – | $7.64 \times 10^{-6}$ | – |

**Table S6.** Network characterization of the tissue structures. Degree and strength ($\sum_j 1/d_{ij}^2$): mean and coefficient of variation; largest connected component (% of cells) and number of components; mean local clustering coefficient; effective dimension from the growth of the number of cells within $k$ steps ($3 \leq k \leq 20$).

| Structure | Cells | $r$ | $\langle k \rangle$ | $CV_k$ | $\langle s \rangle$ | $CV_s$ | LCC (%) | Comp. | $C$ | $d_{\mathrm{eff}}$ |
|---|---|---|---|---|---|---|---|---|---|---|
| lattice | 233,479 | 3 | 6.0 | 0.03 | 0.91 | 0.03 | 100.0 | 1 | 0.40 | 1.87 |
| lattice | 233,479 | 4 | 6.0 | 0.03 | 0.91 | 0.03 | 100.0 | 1 | 0.40 | 1.87 |
| lattice | 233,479 | 6 | 17.9 | 0.04 | 1.43 | 0.03 | 100.0 | 1 | 0.53 | 1.90 |
| lattice | 233,479 | 8 | 35.8 | 0.04 | 1.79 | 0.04 | 100.0 | 1 | 0.56 | 1.90 |
| lattice | 233,479 | 10 | 53.6 | 0.05 | 2.00 | 0.04 | 100.0 | 1 | 0.57 | 1.93 |
| lattice | 233,479 | 12 | 83.2 | 0.05 | 2.24 | 0.04 | 100.0 | 1 | 0.58 | 1.90 |
| section z0 | 233,479 | 4 | 3.5 | 0.89 | 1.12 | 0.93 | 0.5 | 56,105 | 0.53 | 1.24 |
| section z0 | 233,479 | 6 | 7.1 | 0.82 | 1.28 | 0.87 | 3.8 | 13,722 | 0.64 | 1.40 |
| section z0 | 233,479 | 8 | 11.6 | 0.75 | 1.37 | 0.84 | 55.2 | 1,156 | 0.66 | 1.64 |
| section z0 | 233,479 | 10 | 17.0 | 0.69 | 1.44 | 0.82 | 56.3 | 108 | 0.66 | 1.71 |
| section z0 | 233,479 | 12 | 23.5 | 0.64 | 1.49 | 0.80 | 56.3 | 18 | 0.66 | 1.78 |
| section z0 | 233,479 | 15 | 34.7 | 0.58 | 1.56 | 0.77 | 56.3 | 6 | 0.65 | 1.80 |
| section z0 | 233,479 | 20 | 58.5 | 0.51 | 1.63 | 0.75 | 56.3 | 5 | 0.64 | 1.76 |
| section z5 | 193,093 | 4 | 3.3 | 1.03 | 1.09 | 0.99 | 0.5 | 58,787 | 0.48 | 1.19 |
| section z5 | 193,093 | 6 | 6.4 | 0.99 | 1.22 | 0.95 | 1.9 | 20,800 | 0.63 | 1.30 |
| section z5 | 193,093 | 8 | 10.3 | 0.92 | 1.30 | 0.92 | 50.6 | 2,998 | 0.67 | 1.59 |
| section z5 | 193,093 | 10 | 14.7 | 0.85 | 1.35 | 0.90 | 57.4 | 315 | 0.66 | 1.75 |
| section z5 | 193,093 | 12 | 20.6 | 0.78 | 1.40 | 0.89 | 57.7 | 27 | 0.65 | 1.75 |
| section z5 | 193,093 | 15 | 29.9 | 0.71 | 1.45 | 0.87 | 57.7 | 7 | 0.64 | 1.75 |
| section z5 | 193,093 | 20 | 49.5 | 0.61 | 1.52 | 0.84 | 57.7 | 3 | 0.64 | 1.84 |
| section z10 | 228,462 | 4 | 3.7 | 0.93 | 1.21 | 0.93 | 0.7 | 57,199 | 0.52 | 1.47 |
| section z10 | 228,462 | 6 | 7.5 | 0.88 | 1.37 | 0.89 | 4.0 | 16,970 | 0.64 | 1.26 |
| section z10 | 228,462 | 8 | 12.1 | 0.82 | 1.47 | 0.86 | 53.9 | 2,128 | 0.67 | 1.45 |
| section z10 | 228,462 | 10 | 17.9 | 0.77 | 1.54 | 0.84 | 56.9 | 192 | 0.66 | 1.77 |
| section z10 | 228,462 | 12 | 24.4 | 0.73 | 1.60 | 0.83 | 56.9 | 35 | 0.65 | 1.80 |
| section z10 | 228,462 | 15 | 36.5 | 0.67 | 1.67 | 0.81 | 57.0 | 10 | 0.65 | 1.85 |
| section z10 | 228,462 | 20 | 60.5 | 0.60 | 1.75 | 0.78 | 57.0 | 6 | 0.64 | 1.86 |
| section z15 | 154,623 | 4 | 2.7 | 1.05 | 0.98 | 1.00 | 1.1 | 54,477 | 0.46 | 1.22 |
| section z15 | 154,623 | 6 | 5.2 | 1.02 | 1.09 | 0.97 | 2.0 | 23,620 | 0.60 | 1.29 |
| section z15 | 154,623 | 8 | 8.4 | 0.95 | 1.15 | 0.94 | 19.2 | 6,884 | 0.66 | 1.46 |
| section z15 | 154,623 | 10 | 12.3 | 0.88 | 1.20 | 0.92 | 60.1 | 1,394 | 0.67 | 1.78 |
| section z15 | 154,623 | 12 | 16.8 | 0.83 | 1.24 | 0.91 | 60.9 | 227 | 0.66 | 1.76 |
| section z15 | 154,623 | 15 | 25.2 | 0.76 | 1.29 | 0.89 | 60.9 | 26 | 0.65 | 1.82 |
| section z15 | 154,623 | 20 | 41.9 | 0.67 | 1.35 | 0.86 | 60.9 | 8 | 0.64 | 1.79 |
| 3-D tissue | 4,137,043 | 2 | 1.8 | 0.96 | 1.17 | 0.96 | 0.6 | 1,638,058 | 0.26 | 1.51 |
| 3-D tissue | 4,137,043 | 4 | 11.6 | 0.72 | 2.31 | 0.74 | 98.1 | 26,564 | 0.57 | 2.23 |
| 3-D tissue | 4,137,043 | 6 | 33.9 | 0.62 | 3.21 | 0.66 | 99.9 | 696 | 0.55 | 2.01 |
| 3-D tissue | 4,137,043 | 8 | 72.7 | 0.55 | 4.01 | 0.61 | 100.0 | 79 | 0.55 | 1.94 |

**Table S7.** Calibrated z0 points re-simulated on the matched hexagonal lattice (30 replicates each). $J$, fit cost with offsets re-profiled; $f_2$, cells infected or dead at day 2 (%; target $13.0 \pm 5.0$); damage and refractory, permissive cells infected/dead or refractory at day 10 (%); $t_{\mathrm{clear}}$, first day the scaled mean titer falls below the detection limit.

| Calibrated point | Tissue | $J$ | $f_2$ | Peak day | Damage | Refract. | $t_{\mathrm{clear}}$ |
|---|---|---|---|---|---|---|---|
| all four + CTL | section z0 | 43.5 | 9.8 | 4.3 | 58.7 | 0.0 | 8.0 |
| | lattice | 52.1 | 38.4 | 2.6 | 98.2 | 0.0 | 7.7 |
| CTL only | section z0 | 44.5 | 8.4 | 4.3 | 54.5 | 0.0 | 8.0 |
| | lattice | 48.3 | 32.9 | 2.8 | 98.3 | 0.0 | 7.8 |
| BI+RF+BP, no CTL | section z0 | 89.7 | 16.4 | 2.9 | 54.8 | 44.6 | – |
| | lattice | 155.3 | 33.7 | 1.5 | 43.8 | 56.2 | 7.0 |
| none, no CTL (day 6) | section z0 | 105.7 | 17.3 | 3.8 | 97.0 | 0.0 | – |
| | lattice | 66.9 | 57.9 | 2.1 | 100.0 | 0.0 | 8.4 |

**Table S8.** Sensitivity of the three-dimensional tissue to the assumed distance between sections. The twenty sections are stacked 1 unit apart in the reconstruction (an assumption: section thickness and the material lost at each cut are unknown); here the $z$ coordinate is multiplied by the spacing. **Top**, network of a $1200 \times 1200$ window through all sections (single section: z0 in the same window): mean degree, largest connected component, cells within 20 steps ($B_{20}$, 50 start cells at least 250 units from the window edge), and effective dimension. **Bottom**, holes in a $400 \times 400$ window: median size (the radius that bridges a hole), number of holes at least 10, 12, 15, and 20 units in size per $10^4$ cells of one section, and largest hole.

| Spacing | $r$ | $\langle k \rangle$ | LCC (%) | $B_{20}$ | $d_{\text{eff}}$ |
|---|---|---|---|---|---|
| 1 | 4 | 11.9 | 49.5 | 4,802 | 2.26 |
| | 6 | 34.8 | 100.0 | 16,402 | 2.04 |
| | 8 | 74.0 | 100.0 | 31,953 | 1.86 |
| | 10 | 132.4 | 100.0 | 54,606 | 1.90 |
| 2 | 4 | 6.9 | 35.4 | 1,906 | 2.22 |
| | 6 | 20.5 | 99.8 | 13,558 | 2.28 |
| | 8 | 44.1 | 100.0 | 27,253 | 2.13 |
| | 10 | 80.1 | 100.0 | 46,796 | 1.98 |
| 3 | 4 | 5.5 | 19.9 | 2,565 | 1.96 |
| | 6 | 14.5 | 50.0 | 10,588 | 2.45 |
| | 8 | 32.7 | 100.0 | 27,441 | 2.34 |
| | 10 | 59.1 | 100.0 | 44,296 | 2.15 |
| 5 | 4 | 3.5 | 0.4 | 573 | 1.38 |
| | 6 | 10.2 | 48.7 | 5,723 | 2.46 |
| | 8 | 22.0 | 99.9 | 19,211 | 2.44 |
| | 10 | 36.6 | 100.0 | 35,726 | 2.41 |
| single section | 4 | 3.5 | 2.6 | 431 | 1.42 |
| | 6 | 7.0 | 8.9 | 411 | 1.18 |
| | 8 | 11.4 | 50.7 | 909 | 1.61 |
| | 10 | 16.8 | 52.2 | 2,028 | 1.74 |

| Spacing | Median size | $\geq 10$ | $\geq 12$ | $\geq 15$ | $\geq 20$ | Largest |
|---|---|---|---|---|---|---|
| 1 | 4.5 | 78 | 4 | 0 | 0 | 13 |
| 2 | 6.0 | 175 | 27 | 8 | 4 | 21 |
| 3 | 6.3 | 475 | 58 | 12 | 4 | 30 |
| 5 | 8.2 | 2976 | 183 | 27 | 4 | 44 |
| single section | 12.2 | 424 | 291 | 79 | 6 | 28 |

#### 6. Supplementary figures

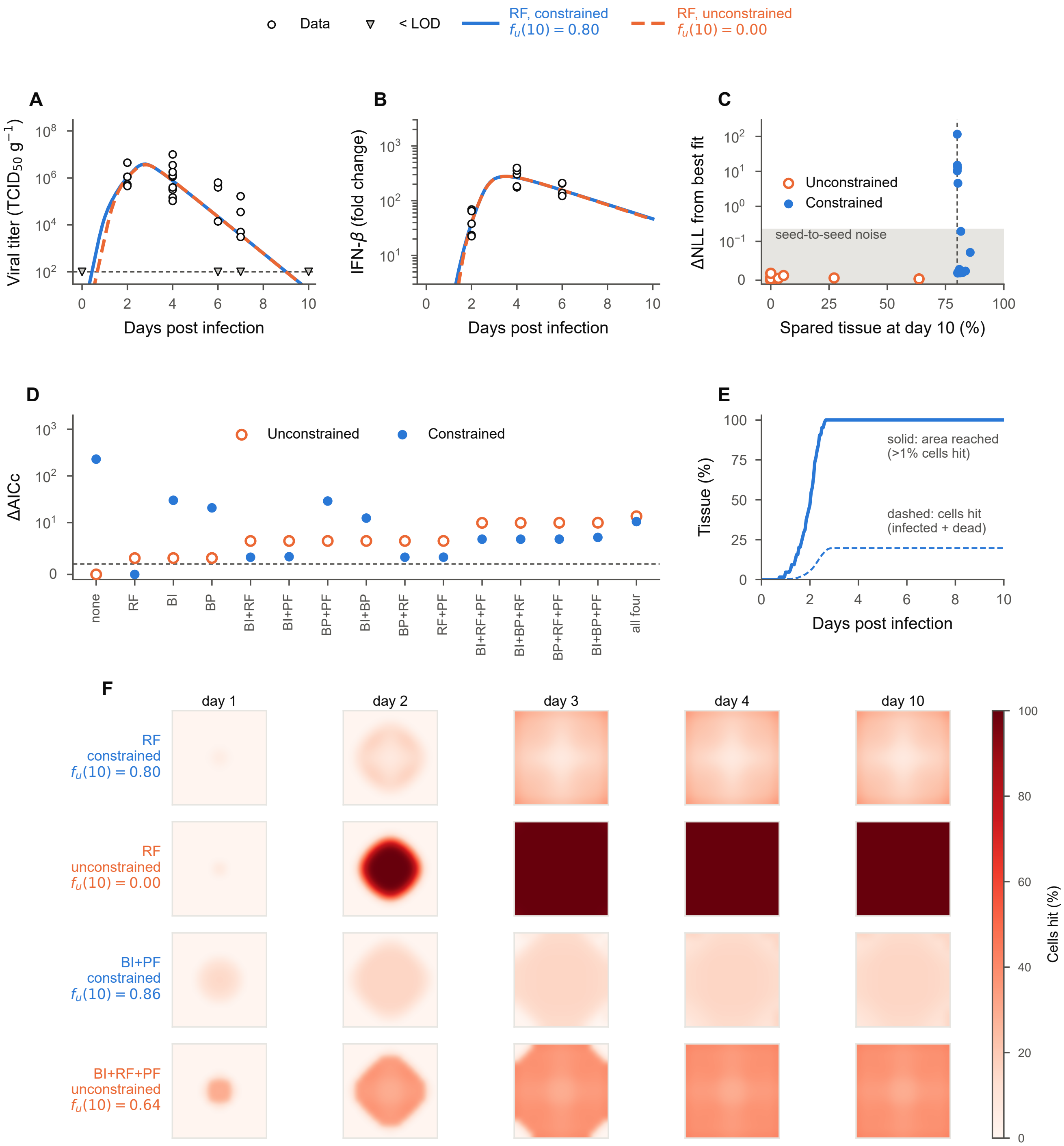


**Fig. S1.** Bulk kinetics do not distinguish IFN mechanisms on homogeneous tissue, and tissue sparing does not produce spatially confined lesions. (*A*,*B*) Experimental viral titers (*A*) and IFN-$\beta$ fold changes (*B*) (open circles; triangles, censored titers; day-0 triangle, inoculum; dashed line, detection limit) and the best-fitting refractory-conversion model with (blue, solid) and without (orange, dashed) the sparing constraint. Model outputs are scaled by profiled offsets. (*C*) Day-10 spared fraction versus loss of fit for 15 mechanism combinations, with (filled) and without (open) the constraint; dashed line, sparing threshold; gray band, variation across optimizer seeds. (*D*) ΔAICc with and without the constraint, ordered by unconstrained ranking; dashed line, $\Delta\text{AICc} = 2$. (*E*) Fraction of the domain reached by infection (solid) and cells infected or dead (dashed) under the constraint. (*F*) Spatial distribution of infected or dead cells at days 1, 2, 3, 4, and 10 for refractory conversion (M05), constrained BI+PF (M10), and unconstrained BI+RF+PF (M14). Row colors indicate constrained (blue) or unconstrained (orange) fits; labels give final spared fractions. Infection reaches the entire domain by day 3, with sparing arising from widespread refractory conversion or uniformly reduced infection rather than spatial confinement.

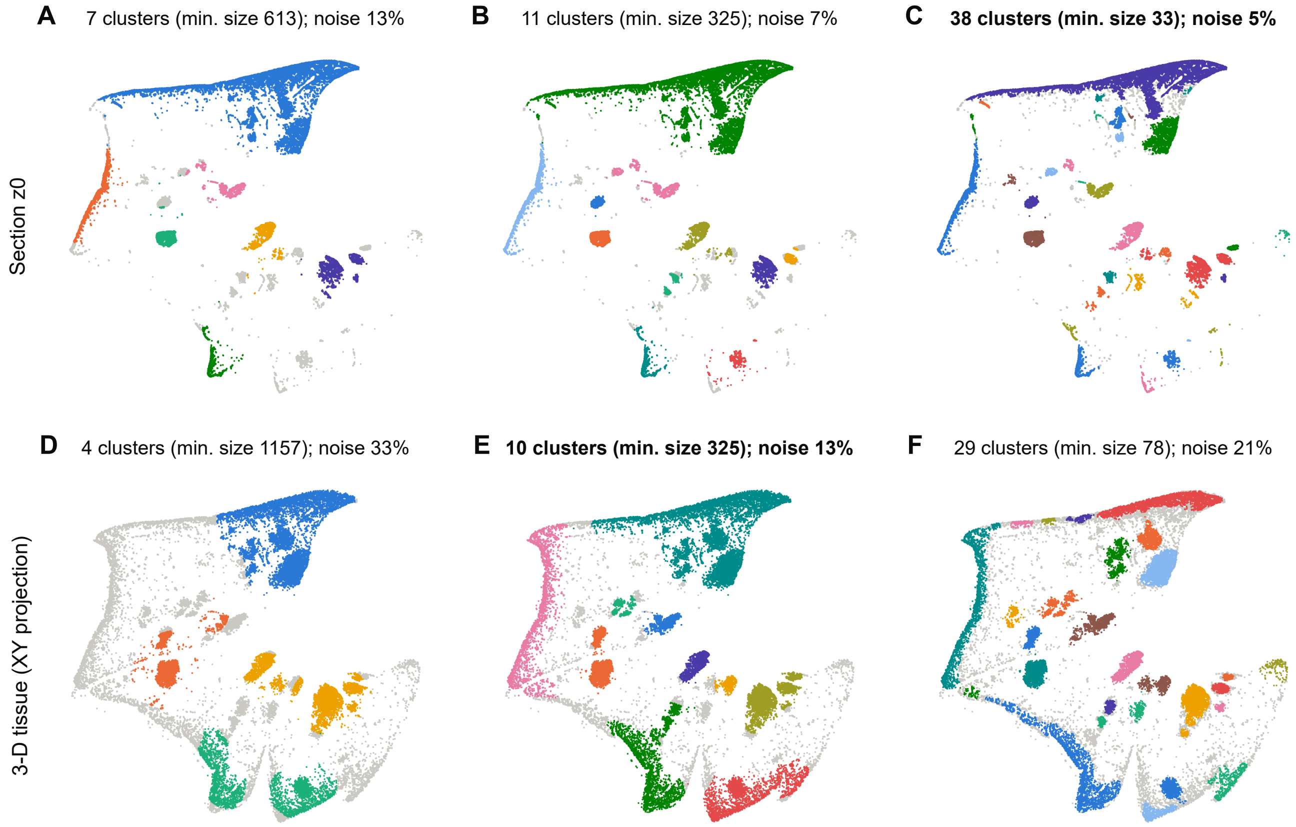


**Fig. S2.** Clusters of IAV-positive cells used to set the inoculum. HDBSCAN applied to the coordinates of the immunostained positive cells, at three values of the minimum cluster size: (*A–C*) the 30,404 positive cells of section z0; (*D–F*) a random subsample of 30,000 positive cells of the three-dimensional tissue, shown in projection. Colors distinguish clusters; gray, cells left unclustered (noise). Bold titles mark the solutions used in the model: 38 clusters in the section ($N_0 = 38$) and 10 in the volume. Coarser settings merge neighboring foci and leave more cells unclustered, finer ones split single foci.

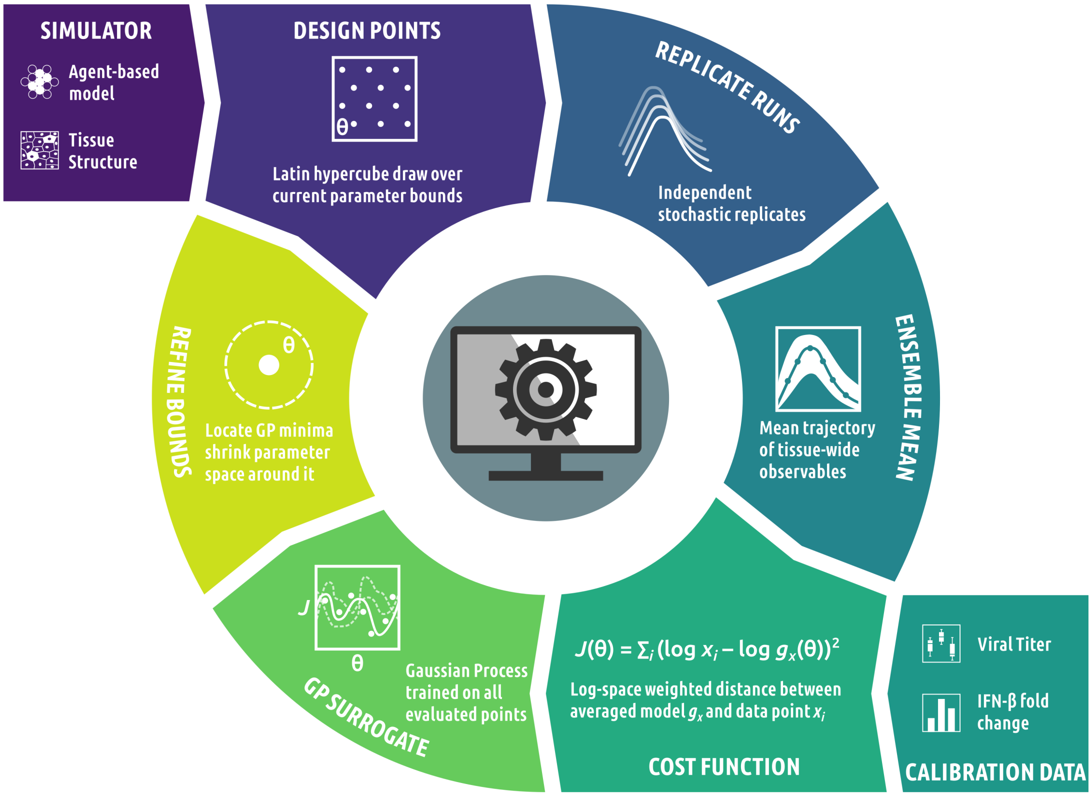


**Fig. S3.** Calibration of the agent-based model. Design points are drawn over the current parameter bounds, each simulated with independent stochastic replicates; the replicate mean of the tissue-wide virus and IFN is compared with the data through the profiled censored likelihood, and the bounds are refined around the best candidates.

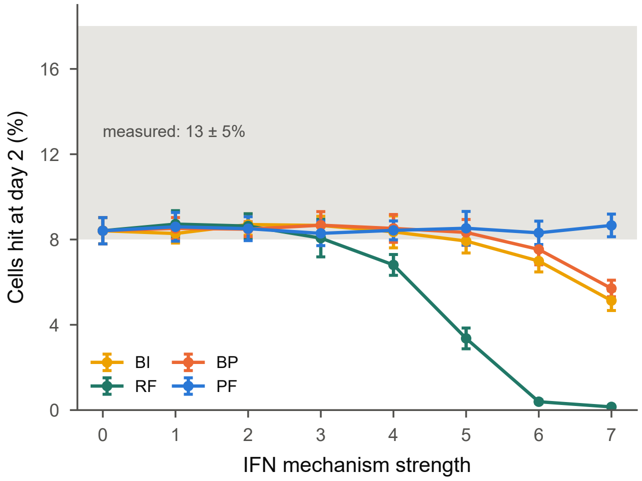


**Fig. S4.** Lesion extent under single IFN mechanisms on section z0. Cells infected or dead at day 2 when each mechanism is added alone to the best z0 model (CTL, no IFN mechanism) at seven strengths (level 0, inert; level 7, strongest; same runs as main-text Fig. 4*A*); mean $\pm$ s.d. of 10 replicates; gray band, measured lesion extent.

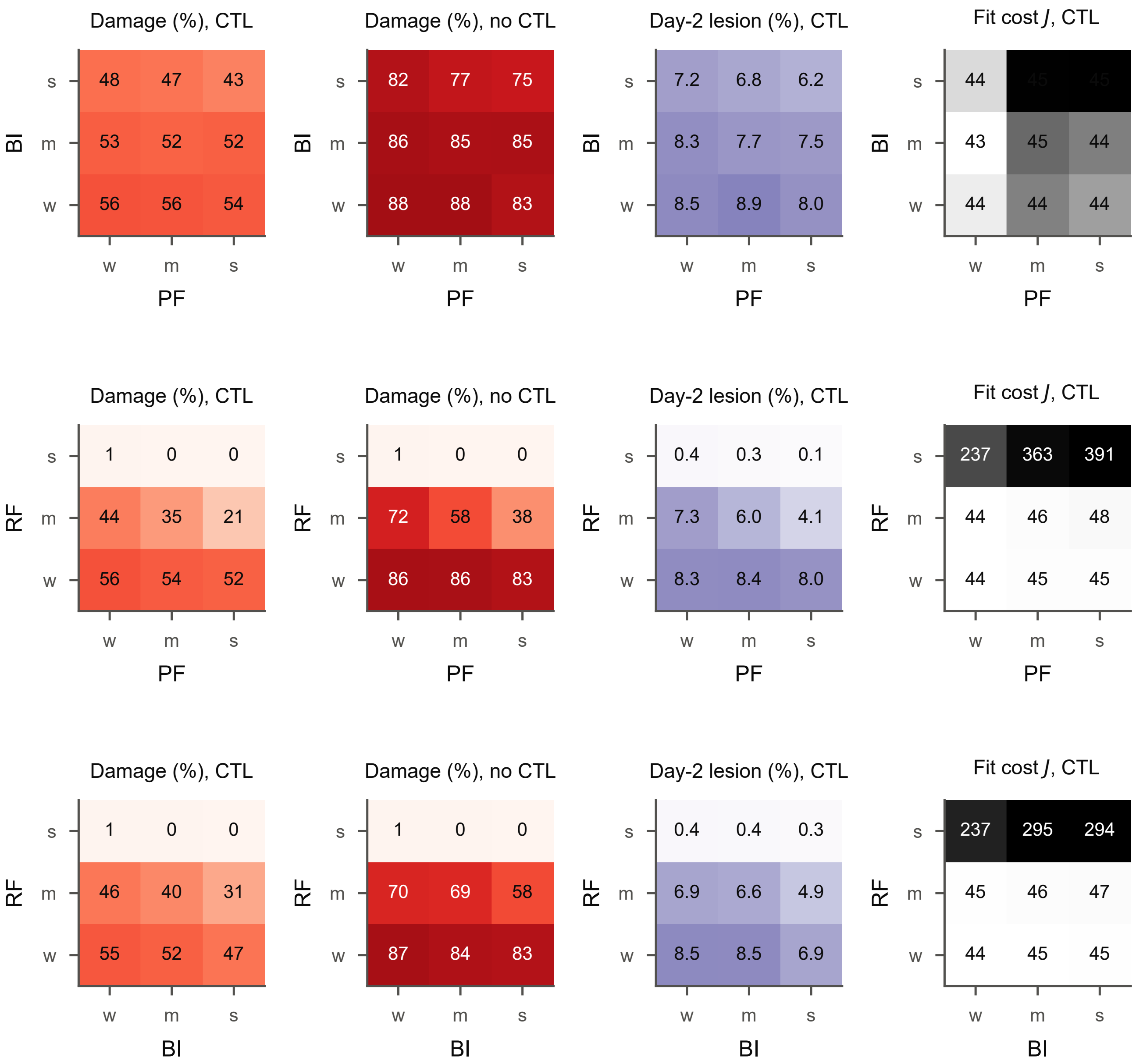


**Fig. S5.** Combinations of IFN mechanisms on section z0. Pairs of mechanisms added to the best z0 model (CTL, no IFN mechanism) at three strengths each (w, weak; m, intermediate; s, strong: $K_{BI} = 8, 4, 2$; $K_R = 200, 50, 12$; $K_{PF} = 10^7, 10^5, 10^3$ with $\alpha_{PF} = 10$), 8 replicates per cell. Columns: permissive cells infected or dead at day 10 with and without CTL; cells infected or dead at day 2 with CTL; fit cost to the viral and IFN data with CTL. Positive feedback reinforces refractory conversion but barely changes blocking of infection.

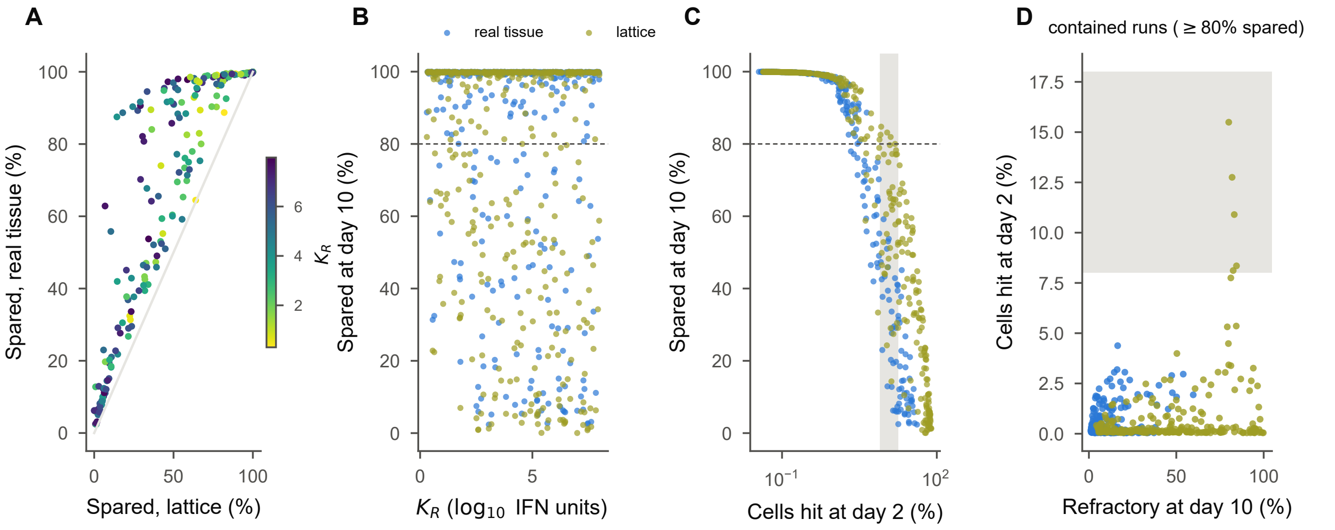


**Fig. S6.** IFN-only containment on real and homogeneous tissue. 400 Latin-hypercube sets of IFN-mechanism and transport parameters (all four mechanisms on, CTL off), each simulated on section z0 and on the matched lattice (2 replicates). (*A*) Spared permissive cells at day 10 on both tissues, colored by the refractory half-maximum $K_R$. (*B*) Spared cells against $K_R$. (*C*) Spared cells against the day-2 lesion extent; gray band, measured. (*D*) Among contained runs ($\geq 80\%$ spared), refractory fraction at day 10 against the day-2 lesion: on the section containment is abortive; on the lattice it proceeds by tissue-wide refractory conversion, the only route to a lesion of the measured size.

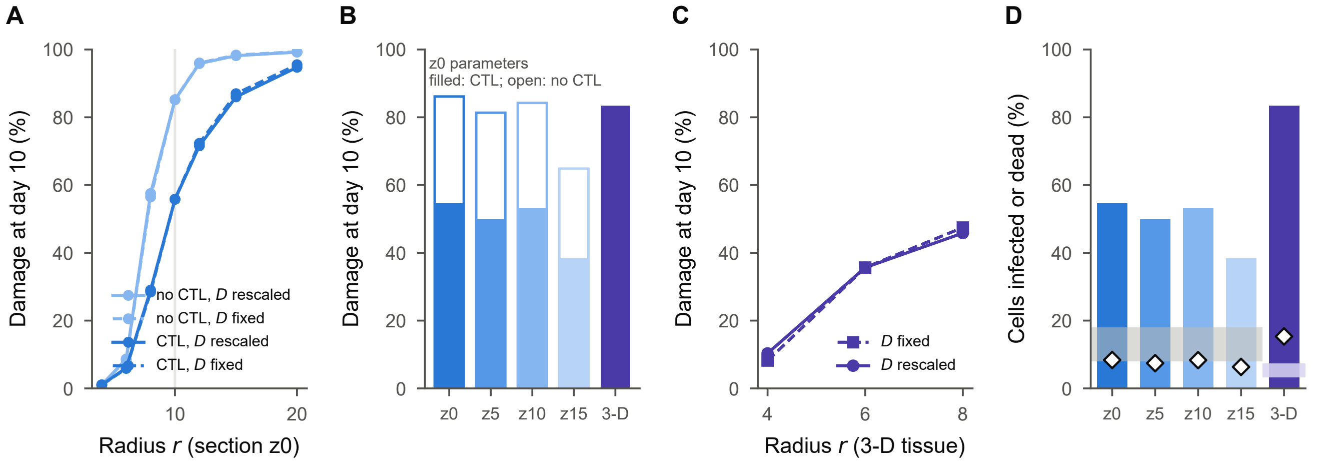


**Fig. S7.** Spread depends on the neighborhood radius, not on the coupling strength. (*A*) Damage at day 10 for the best z0 model at radii from 4 to 20, with diffusivities fixed (dashed) or rescaled to keep the mean coupling constant (solid), with and without CTL; the gray line marks the calibrated radius. (*B*) Damage at day 10 for the same parameters on four sections and on the 3-D tissue (filled, CTL; open, no CTL). (*C*) Damage at day 10 for the best 3-D model at $r = 4$, 6, and 8, with fixed or rescaled diffusivities. (*D*) The z0 parameters with CTL on four sections and on the 3-D tissue: damage at day 10 (bars) and cells infected or dead at day 2 (diamonds); gray band, measured lesion extent in z0; purple band, measured over the volume.

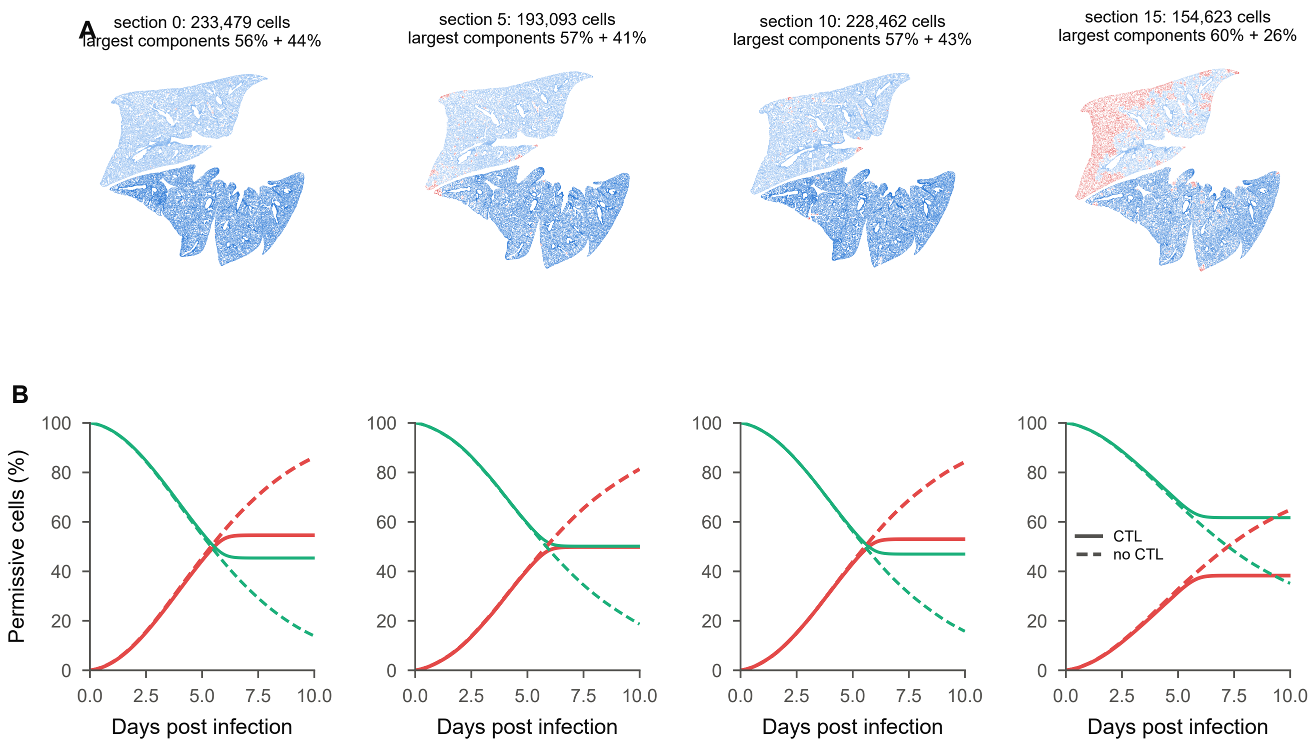


**Fig. S8.** Other sections of the same lung. (*A*) Sections 0, 5, 10, and 15 of the reconstructed volume colored by connected component at $r = 10$ (dark and light blue, the two largest components; red, smaller components). (*B*) Permissive cells infected or dead (red) and susceptible (green) over time for the best z0 model simulated unchanged on each section with CTL (solid) and without (dashed); mean of 50 (z0) or 10 replicates. The least connected section (z15, mean degree 12.3 versus 17.0 in z0) loses the fewest cells.

#### 7. Supplementary movies

**Movie S1.** Infection on the matched hexagonal lattice (file Mov_S1.gif).
The best z0 model (CTL, no IFN mechanism) simulated on the hexagonal lattice matched to section z0 in number of cells, mean degree, and mean coupling strength (233,479 cells, radius 6); same run as the lattice row of main-text Fig. 2*D*, one replicate, frames every 12 h from day 0 to day 10. Left, cell states (green, healthy, i.e., susceptible or non-permissive; red, infected; dark gray, dead); middle, extracellular virions per cell; right, IFN per cell (arbitrary units), both on logarithmic color scales. For display the lattice is binned on a regular grid and colors are averaged within each bin. Foci grow as circles, merge within the first days, and consume nearly all permissive cells before CTL become effective.

**Movie S2.** Infection on section z0 (file Mov_S2.gif).
The same model and parameters on the reconstructed section z0; same run as the real-tissue row of main-text Fig. 2*D*, one replicate, frames every 12 h from day 0 to day 10. Panels and color scales as in Movie S1. Foci grow irregularly along the tissue network, remain separated by airspaces and sparse tissue, and the virus is cleared after CTL expand, leaving about half of the permissive cells intact.

1. Esteban A Hernandez-Vargas, Esther Wilk, Laetitia Canini, Franklin R Toapanta, Sebastian C Binder, Alexey Uvarovskii, Ted M Ross, Carlos A Guzmán, Alan S Perelson, and Michael Meyer-Hermann. Effects of aging on influenza virus infection dynamics. *Journal of Virology*, 88(8):4123–4131, 2014.
2. Timo Frensing, Sascha Y Kupke, Mandy Bachmann, Susanne Fritzsche, Lili E Gallo-Ramirez, and Udo Reichl. Influenza virus intracellular replication dynamics, release kinetics, and particle morphology during propagation in mdck cells. *Applied Microbiology and Biotechnology*, 100(16):7181–7192, 2016.
3. The GUDHI Project. *GUDHI User and Reference Manual*. GUDHI Editorial Board, 3.13.0 edition, 2026.
4. Pavitra Roychoudhury, David A Swan, Elizabeth Duke, Lawrence Corey, Jia Zhu, Veronica Davé, Laura Richert-Spuhler, Jennifer M. Lund, Martin Prlic, and Joshua T Schiffer. Tissue-resident T cell–derived cytokines eliminate herpes simplex virus-2–infected cells. *The Journal of Clinical Investigation*, 130(6):2903–2919, 2020. doi: 10.1172/JCI132583.
5. Melanie E. Moses, Steven Hofmeyr, Judy L. Cannon, Akil Andrews, Rebekah Gridley, Monica Hinga, Kirtus Leyba, Abigail Pribisova, Vanessa Surjadidjaja, Humayra Tasnim, and Stephanie Forrest. Spatially distributed infection increases viral load in a computational model of SARS-CoV-2 lung infection. *PLOS Computational Biology*, 17(12):e1009735, December 2021. doi: 10.1371/journal.pcbi.1009735.
6. Hanyu Li, Kazuki Kuga, and Kazuhide Ito. Visual prediction and parameter optimization of viral dynamics in the mucus milieu of the upper airway based on CFPD-HCD analysis; viral dynamics and parameterization. *Computer Methods and Programs in Biomedicine*, 238:107622, 2023. doi: 10.1016/j.cmpb.2023.107622. Epub 2023-05-25 (per PubMed).
7. Benjamin Raach, Nils Bundgaard, Marika J. Haase, Jörn Starruß, Rocio Sotillo, Megan L. Stanifer, and Frederik Graw. Influence of cell type specific infectivity and tissue composition on SARS-CoV-2 infection dynamics within human airway epithelium. *PLOS Computational Biology*, 19(8):e1011356, August 2023. doi: 10.1371/journal.pcbi.1011356.
8. Armand Bankhead, Emiliano Mancini, Amy C. Sims, Ralph S. Baric, Shannon McWeeney, and Peter M. A. Sloot. A simulation framework to investigate in vitro viral infection dynamics. *Journal of Computational Science*, 4(3):127–134, May 2013. doi: 10.1016/j.jocs.2011.08.007.
9. Josua O. Aponte-Serrano, Jordan J. A. Weaver, T. J. Sego, James A. Glazier, and Jason E. Shoemaker. Multicellular spatial model of RNA virus replication and interferon responses reveals factors controlling plaque growth dynamics. *PLOS Computational Biology*, 17(10):e1008874, October 2021. doi: 10.1371/journal.pcbi.1008874.
10. Catherine Beauchemin, John Samuel, and Jack Tuszynski. A simple cellular automaton model for influenza A viral infections. *Journal of Theoretical Biology*, 232(2):223–234, 2005. doi: 10.1016/j.jtbi.2004.08.001.
11. T. J. Sego, Ericka D. Mochan, G. Bard Ermentrout, and James A. Glazier. A multiscale multicellular spatiotemporal model of local influenza infection and immune response. *Journal of Theoretical Biology*, 532:110918, 2022. doi: 10.1016/j.jtbi.2021.110918.
12. Sadegh Marzban, Renji Han, Nóra Juhász, and Gergely Röst. A hybrid PDE–ABM model for viral dynamics with application to SARS-CoV-2 and influenza. *Royal Society Open Science*, 8(11): 210787, 2021. doi: 10.1098/rsos.210787.
13. Alexey Tokarev, Anastasia Mozokhina, and Vitaly Volpert. Competition of SARS-CoV-2 variants in cell culture and tissue: Wins the fastest viral autowave. *Vaccines*, 10(7):995, 2022. doi: 10.3390/vaccines10070995.
14. Ferenc A. Bartha, Nóra Juhász, Sadegh Marzban, Renji Han, and Gergely Röst. In silico evaluation of Paxlovid's pharmacometrics for SARS-CoV-2: A multiscale approach. *Viruses*, 14(5):1103, 2022. doi: 10.3390/v14051103.
15. Christian Quirouette, Nada P Younis, Micaela B Reddy, and Catherine AA Beauchemin. A mathematical model describing the localization and spread of influenza A virus infection within the human respiratory tract. *PLoS Computational Biology*, 16(4):e1007705, 2020.
16. Alex Chen, Timothy Wessler, and M. Gregory Forest. Antibody protection from SARS-CoV-2 respiratory tract exposure and infection. *Journal of Theoretical Biology*, 557:111334, 2023. doi: 10.1016/j.jtbi.2022.111334.
17. Catherine Beauchemin. Probing the effects of the well-mixed assumption on viral infection dynamics. *Journal of Theoretical Biology*, 242(2):464–477, 2006. doi: 10.1016/j.jtbi.2006.03.014.
18. Hugh Mitchell, Drew Levin, Stephanie Forrest, Catherine A. A. Beauchemin, J. Tipper, J. Knight, N. Donart, R. Colby Layton, J. Pyles, P. Gao, Kevin S. Harrod, Alan S. Perelson, and Frederick Koster. Higher Level of Replication Efficiency of 2009 (H1N1) Pandemic Influenza Virus than Those of Seasonal and Avian Strains: Kinetics from Epithelial Cell Culture and Computational Modeling. *Journal of Virology*, 85(2):1125–1135, 2011. doi: 10.1128/JVI.01722-10.
19. G Michael Lavigne, Hayley Russell, Barbara Sherry, and Ruian Ke. Autocrine and paracrine interferon signalling as 'ring vaccination'and 'contact tracing'strategies to suppress virus infection in a host. *Proceedings of the Royal Society B*, 288(1945):20203002, 2021.
20. John Whitman, Aleya Dhanji, Fernand Hayot, Stuart C Sealfon, and Ciriyam Jayaprakash. Spatio-temporal dynamics of host-virus competition: a model study of influenza a. *Journal of Theoretical Biology*, 484:110026, 2020.
21. Yimei Li, Bjarke Frost Nielsen, Simon A Levin, Aartjan JW Te Velthuis, and Bryan T Grenfell. Spatio–temporal modelling of in vitro influenza a virus infection: The impact of defective interfering particles on the type i interferon response. *PLoS Computational Biology*, 22(4):e1014198, 2026.
22. Ulrik Hvid, Bjarke Frost Nielsen, and Kim Sneppen. Bridging spatial scales in respiratory viral infection. *Proceedings of the National Academy of Sciences*, 123(43):e2614436123, 2026. doi: 10.1073/pnas.2614436123.
23. Tom J Howat, Cristina Barreca, Peter O'Hare, Julia R Gog, and Bryan T Grenfell. Modelling dynamics of the type I interferon response to in vitro viral infection. *Journal of The Royal Society Interface*, 3(10):699–709, 2006. doi: 10.1098/rsif.2006.0136.
