## Supplementary figures and images for "Lung tissue organization shapes influenza spread and immune control"

### Supplemental Movie 1

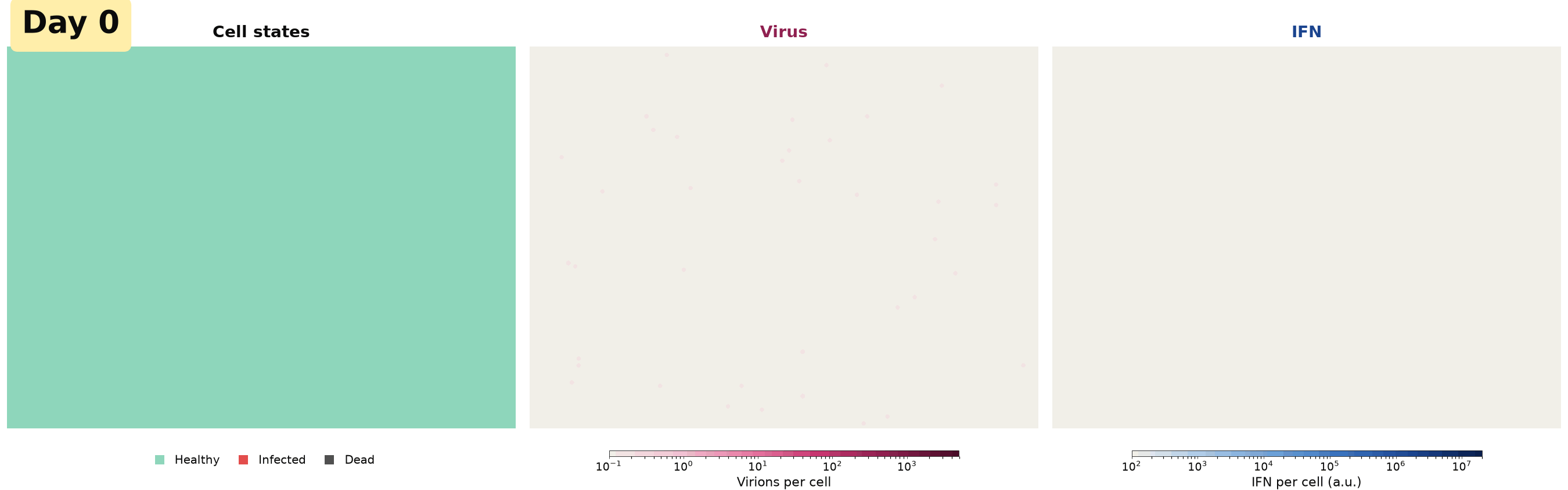

### Supplemental Movie 2

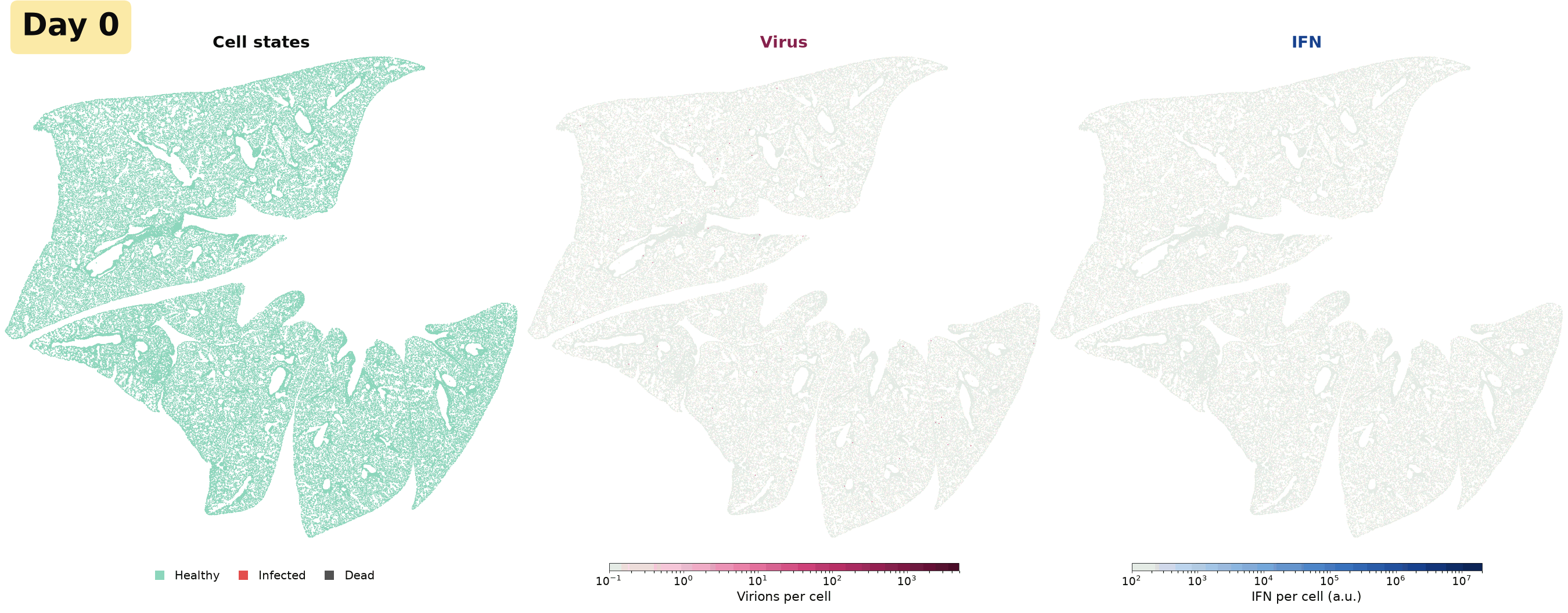
